# Impact of heat and sex on embryonic brain growth in dragons with temperature induced sex-reversal (Pogona vitticeps, Lacertilia: Agamidae)

**DOI:** 10.64898/2026.10.26.761761

**Authors:** Daniel Hoops, Annalise D. Ceddia, Richard Peters, Sarah Whiteley, Shaun P. Collin, Jenna M. Crowe-Riddell

## Abstract

Climate warming can influence sex determination and embryonic growth in oviparous reptiles that rely on external temperatures during development. Warmer incubation temperatures can stimulate growth in ectotherms until oxygen supply can no longer keep pace with growth rate; at which point metabolically expensive organs like the brain suffer. We aimed to understand the impact of heat and sex on brain development in central bearded dragons, *Pogona vitticeps* (Agamidae), a uniquely thermosensitive species that exhibits sex-reversal at warm incubation temperatures. Three sexual phenotypes are possible in *P. vitticeps*: genetically driven ZW females and ZZ males, with the latter phenotype thermally overridden at incubation temperatures >32 °C to induce ZZ sex-reversed females. To visualise changes in brain anatomy, we used buffered Lugol’s iodine to perform contrast-enhanced micro computed tomography (µCT) and synchrotron propagation-based phase-contrast CT scanning. A total of 57 embryos were sampled across developmental stages and incubated at two temperature treatments: cool (28 °C, *n* = 26) and warm (36 °C, *n* = 21). Using high-resolution models of neuroanatomy, we found that the telencephalic region of the forebrain is significantly larger in individuals with ZZ genotypes (males and sex-reversed females) compared to concordant ZW females. Although incubation temperature significantly influenced growth rate and body size during development, overall brain size was not significantly different across temperature treatments. Our results indicate that the effects of genotypic sex on brain development are more important than thermal influences, and thus the brain may be somewhat buffered to the effects of warm incubation temperatures in this arid-adapted dragon.

## Introduction

Anthropogenic climate change poses unique challenges to ectothermic reptiles because they rely on environmental temperatures for physiological, developmental, and behavioural processes [Huey, and Pianka, 1983; Angilletta, 2009]. Although adult ectotherms can often cope with rising temperatures by thermoregulating, i.e. modifying behaviour to keep stable body temperatures [Black, and Tattersall, 2017], embryos of oviparous lizards are more susceptible to heat stress. One potential impact of heat stress is on brain development, which can have flow-on effects for cognition and behaviour into adulthood [Beltrán et al., 2021; Siviter et al., 2017a]. However, the exact relationship between heat stress, the brain, and behaviour is unclear, and the precise effects of rising temperatures appear to be taxon-dependent, likely due to a dearth of data among thermosensitive taxonomic groups [Beltrán et al., 2021].

Temperature-dependent sex determination, where incubation temperature determines the sex of the offspring, is common in reptiles [Capel, 2017]. Sex-reversal is rarer, occurring when the genetic sex determination is overridden or “reversed” by certain egg incubation temperatures [Holleley et al., 2016]. The central bearded dragon, *Pogona vitticeps* (Agamidae), is one of only two squamate species known to reverse their genetically-determined sex in response to hotter incubation temperatures [Holleley et al., 2015; Whiteley et al., 2021; Whiteley et al., 2017]; the other species being the Eastern three lined skink, *Acritoscincus duperreyi* (Scincidae) [Radder et al., 2008; Shine et al., 2002]. Central bearded dragons are oviparous lizards that live in the Australian arid zone and whose genetic sex determination results in heterogametic females (ZW sex chromosomes) and homogametic males (ZZ sex chromosomes). In *P. vitticeps*, incubation temperatures hotter than 32 °C can cause complete feminisation of a ZZ embryo, which then develops into a reproductively viable female with functional ovaries and oviducts [Holleley et al., 2015]. These sex-reversed females (ZZf) show a unique set of characteristics not seen in their concordant male (ZZm) and female (ZWf) counterparts [Li et al., 2016]. ZZf hatchlings tend to have larger relative tail lengths and body masses and exhibit bolder and more active behaviours [Li et al., 2016]. Thermosensitive species like *P. vitticeps*, therefore, provide useful models for exploring the intertwined influences of genetic and environmental effects on sex, brain, and behaviour.

a Heat stress can cause detrimental effects on brain development. The physiological mechanism is thought to be an increased demand for cellular oxygen at higher metabolic rates, which ultimately limits the rate of cell proliferation and/or shuts down spontaneous neural activity [Andreassen et al., 2022; Stepien, and Wielockx, 2024]. The brain is particularly energetically expensive to build and maintain and therefore may be particularly susceptible to heat-induced oxygen deprivation (i.e. an evolutionary developmental extension of the expensive brain hypothesis) [Laughlin, 2001; Song et al., 2023]. Furthermore, we might expect oxygen deficiency to have a heterogeneous effect across the brain depending on the energetic requirements associated with the development of different brain regions (i.e. an evolutionary developmental extension of the mosaic brain evolution hypothesis) [Barton, and Harvey, 2000]. Therefore, the effects of warm incubation temperatures may be specific to certain brain regions and their specific functions.

Mosaic patterns of brain evolution may reflect differences in the timing of neural development, known as the “late equals large” theory [Finlay, and Darlington, 1995; Finlay et al., 2001]. In mammals neurogenesis occurs sequentially from posterior to anterior brain regions, and late-developing forebrain structures, such as the telencephalon, are especially susceptible to developmental influences and are relatively large compared to the rest of the brain [Finlay, and Darlington, 1995; Finlay et al., 2010]. Furthermore, higher incubation temperatures have been linked to deficits in social learning in *P. vitticeps* that lasted into adulthood [Siviter et al., 2017a]; these brain functions are largely mediated by the telencephalon. We are therefore particularly interested in whether the telencephalon is specifically affected by high incubation temperatures.

To explore the entwinned influences of heat and sex on brain development, we examined brain anatomy in *P. vitticeps* embryos that were exposed to warm and cool incubation temperatures. High resolution models of the telencephalon and whole brain were reconstructed from contrast-enhanced bioimaging from early development to pre-hatchling. We compared brain and telencephalon sizes between males (ZZm) and females (ZWf) incubated at normal temperatures (28 °C), and males (ZZm) and sex-reversed females (ZZf) produced at hotter temperatures (36 °C). We predicted that hot-incubated embryos (ZZm and ZZf) would have smaller brains and/or telencephala compared to cool-incubated embryos (ZZm, ZWf) due to the purported neural cost associated with higher metabolic rate/oxygen deprivation induced by heat stress during development.

## Methods

### Specimens

The embryos we used were collected and analysed as part of a previous study and the details of their breeding and collection are described in Whiteley et al. [2017]. Briefly, eggs were split between a cool incubation temperature (28°C), for normal embryonic development, or a warm incubation temperature (36°C), to induce sex-reversal (Figure 1a). In both treatments, all embryos with a ZW genotype develop as females (ZWf/concordant females). In the cool treatment, embryos with a ZZ genotype develop as males (ZZm/concordant males), whereas in the warm treatment embryos with a ZZ genotype are more likely to develop as females (ZZf/sex-reversed females). Embryo genotype and sex were previously determined by a PCR-based test and histology of the gonadsas part of the study by Whiteley et al [2017].

**Figure 1.**
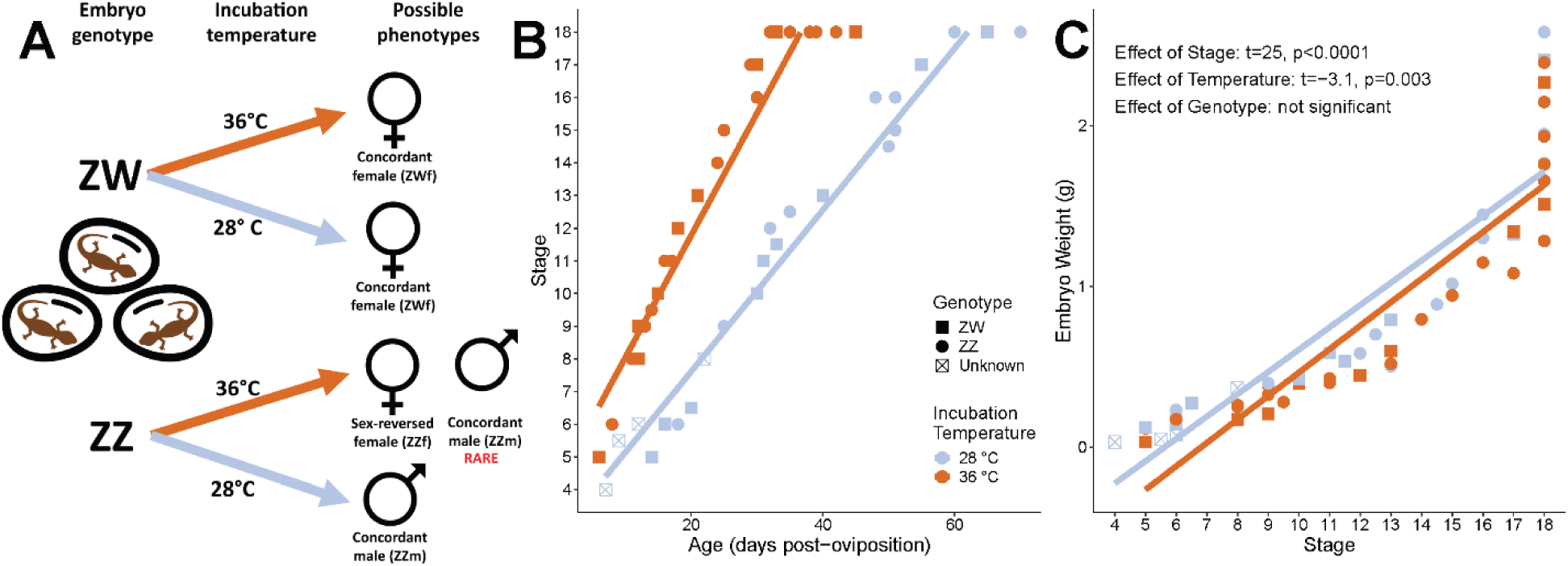
Development of central bearded dragon (*Pogona vitticeps)* embryos incubated at two different temperatures. **A.** Bearded dragon embryos inherit either ZW or ZZ sex chromosomes and, when incubated at a relatively cool temperature (28°C, blue), develop into the sex dictated by their genotype: concordant females (ZWf) and concordant males (ZZm). When incubated at a relatively warm temperature (36°C, orange), ZZ-genotype embryos usually develop as sex-reversed females (ZZf). Rarely, 36°C, ZZ-genotype embryos may develop into concordant males (ZZm). The 36°C, ZW-genotype embryos develop as concordant females (ZWf). **B.** Embryos incubated at the warmer temperature develop faster, progressing more rapidly through developmental stages. For accurate comparisons, stage, rather than age, is used as the predictor variable when comparing across incubation temperatures. **C.** Embryos incubated at the warmer temperature weigh less than embryos incubated at the cooler temperature. Genotype does not predict embryo weight.

Across development, embryos were manually extracted from their eggs and immersed in 10% neutral-buffered formalin for 24-72 hours, then rinsed in water and stored in 70% ethanol-in-water [Whiteley et al., 2017]. After fixation each embryo was weighed and assigned a developmental stage. Developmental stages were designated by Whiteley et al. [2017] based on developmental milestones such as limb and gonad development, resulting in 18 distinct stages of embryonic development.

### Iodine staining and bioimaging

For bioimaging, all embryos were incrementally rehydrated in decreasing ethanol concentrations (70% to 50% to 25%) then immersed in 1.25% Sorensen’s-buffered Lugol’s iodine solution (pH 7.2) for 6 to 55 hours following an established staining protocol [Metscher, 2009a; Dawood et al., 2021]. Specimens were imaged at The University of Melbourne’s Trace Analysis for Chemical, Earth and Environmental Sciences platform (TrACEES; 48 specimens) or at the Australian Synchrotron’s Imaging and Medical Beamline (IMBL; 9 specimens). After imaging, embryos were dehydrated and leached of Lugol’s iodine stain in increasing ethanol concentrations (25% to 50% to 70%) until the ethanol remained clear and the specimen returned to pre-stain hue.

At the TrACEES facility, specimens were imaged by micro computed tomography (µCT) using a Phoenix Nanotom M system (Waygate Technologies), operated using XS Control and Phoenix Datos|x acquisition and reconstruction software. Specimens were individually transferred into microcentrifuge tubes containing Sorensen’s buffer and suspended between the X-ray source and detector to keep their heads in the full field of view and to allow an optimised voxel resolution, which ranged from 2.75 to 9 µm. Specimens were scanned for 10 min with an

X-ray tube voltage of 120 kV and a current of 300 μA, collecting 1199 2D X-ray projections from different angles through a full 360-degree rotation of the sample. An in-line median filter, a region-of-interest filter, and a beam hardening correction filter were applied during sample reconstruction using the Datos|x reconstruction software to reduce noise.

At the Australian Synchrotron, specimens were imaged with propagation-based phase-contrast computed tomography (PB-CT/PCT) by the Australian Nuclear Science and Technology Organization using their superconducting multipole wiggler-based hard X-ray beamline [Arhatari et al., 2021]. Specimens were individually wrapped in plastic wrap then transferred into microcentrifuge tubes, mounted with a source-to-sample distance of 130 mm and a sample to detector distance of 50 mm. Scanning took 13 minutes with a photon energy of 35 kEv collecting 2640 projections from different angles through a full 360-degree rotation of the sample. The resulting images had an isotropic resolution of 13.3 µm.

### Developmental Staging

Reconstructed image files were imported into and processed using Dragonfly 3D World (Comet Technologies Canada). The differentiation of the neural tube into distinguishable brain regions was then visually assessed for each stage, and the resulting observations recorded as a staging guide for brain development in *Pogona* embryos.

### Digital segmentation and cleaning of 3D brain models

The brain of each specimen was digitally isolated (i.e. segmented) from the rest of the tissue using the Segmentation menu in Dragonfly. To ensure brain models were comparable across scans, the olfactory bulbs, olfactory tracts, optic nerve, spinal cord, and cranial nerves were not included. In particular, we excluded the olfactory bulbs and tracts because these small, thin structures could not be reliably segmented due to differences in staining and scanning quality. Segmentation boundaries for the brain and telencephalon were determined using brain atlases for the bearded dragon and another amphibolurine agamid lizard [Foss et al., 2022; Hoops et al., 2018].

All segmentations (whole brains and telencephala) were converted into contour meshes and exported as digital surface models. Brain meshes were simplified, cleaned, and smoothed in MeshLab [Cignoni et al., 2008]. Surface decimation was used to reduce the total number of faces of each mesh to the lowest number feasible to retain the shape and texture present in the CT scans. The simplified surface meshes had 100 000 and 24 000 faces for the brain and telencephalon, respectively.

To obtain volumes, the cleaned and smoothed meshes were re-imported into Dragonfly, which automatically lists the volumes of each object. To obtain linear measurements, the whole brain meshes were imported into imageJ. The imageJ measuring tool was used to obtain eight measurements for each brain (Supplementary Figure 1).

### Statistical analyses

To analyze brain sizes, the mass of the brain and telencephalon were calculated as the product of their respective volumes and the density of reptile brain tissue, i.e.1.036 g/mL [Font et al., 2019]. The “rest of the brain” was calculated as overall brain mass minus the mass of the telencephalon. Weights were transformed using the natural logarithm and then centred and scaled prior to analysis. Developmental stage was also centred and scaled for analysis.

Linear models analyzing weights were conducted using the stats package in the R statistical computing environment [R Core Team, 2026]. We used incubation temperature (28°C or 36°C), either genotype (ZW or ZZ) or phenotype (ZWf, ZZm, or ZZf), and developmental stage as predictor variables. Developmental stage was used in place of chronological age because, as has been reported previously[Whiteley et al., 2017], warm-incubated embryos develop faster than cool-incubated embryos (Figure 1b). To examine differences in overall embryo growth, we used whole embryo weight as the response variable. To examine differences in brain growth, we used telencephalon or rest-of-the-brain weight as the response variable. To examine differences in allometry, we conducted the same analyses with embryo weight substituted for developmental stage as a predictor.

To analyse our linear measurements of brain shape, we first transformed our data using the log-shape ratio [Mosimann, 1970; Klingenberg, 2016]. We then modelled our data using a permutational ANOVA [Anderson, 2001; Goodall, 1991; Baken et al., 2021]. We used incubation temperature (28°C or 36°C), either genotype (ZW or ZZ) or phenotype (ZWf, ZZm, or ZZf), and developmental stage as predictor variables. We then reduced dimensionality and visualised shape differences between groups using a principal components analysis.

The geometric means calculated by the log-shape ratio transformation are an alternative measure of size to weights. We used this measure as a conceptual replicate of the weight analyses we describe above. We used ANOVAs with incubation temperature (28°C or 36°C), either genotype (ZW or ZZ) or phenotype (ZWf, ZZm, or ZZf), and developmental stage (4–18) as predictor variables and geometric mean as the response variable.

Our shape analyses were conducted primarily using the R package geomorph [Baken et al., 2021], supplemented with the stats [R Core Team, 2026] and car [Fox, and Weisberg, 2019] packages.

During our analysis development process, we generated linear models as above but including all two-way interactions between predictor variables. In all cases, the interaction terms were not significant. Therefore, we present results from models with only main effects included.

Graphs were generated primarily using the R package ggplot2 [Wickham, 2016], supplemented with functions from the R packages cowplot [Wilke, 2015], ggeffects [Lüdecke, 2018], ggforce [Pedersen, 2016], and microViz [Barnett et al., 2021]. The colours used for graphing data were derived from a palette developed by the authors from the colouration of bearded dragons, both wild-type and colour mutations found in the pet trade.

### Data and Code Availability

The complete R code, summary statistics, and model outputs for every analysis are presented in our Statistics Supplement (Supplementary Material 1). Additional analyses are presented in our Supplementary Figures & Tables document (Supplementary Material 2).

All CT scans used in this study are freely available as reconstructed tomographic stacks of tiff-format images via FigShare (https://doi.org/10.25909/c.8789142). Each download folder also contains a text file with the image resolution, a Dragonfly-format file containing the CT scan and its associated segmentations (telencephalon and whole brain) and STL format surface meshes of the telencephalon and whole brain.

## Results

### Embryonic development

Embryonic weight was affected by incubation temperature, with cool-incubated lizards weighing more than warm-incubated lizards (Figure 1c; Statistics Supplement). However, genotype did not predict embryo weight. These results are consistent with a reanalysis of the complete data set of embryo weights from Whiteley et al. [2017] (Supplementary Figure 2). The sexual phenotype of the embryo also did not predict embryo weight (Supplementary Figure 3).

### Brain staging

We characterised the gross morphological changes of the brain from stage 4 until stage 18, the final stage before hatching [Whiteley et al., 2017]. Table 1 outlines the changes we observe qualitatively, excluding stage 7, for which we did not have specimens available. Figure 2 illustrates the major brain regions (and associated structures) visible on the exterior of the brain that were used to define the morphology of the brain at each developmental stage. Figure 3 visually maps the development of the brain across stages. Brains at early embryonic stages (4-10) undergo prominent morphological changes, making it feasible to determine the stage of an embryo based on the shape of its brain. Brains at later embryonic stages (11-18) demonstrate fewer gross changes, with remaining development mostly characterised by changes in size rather than shape.

**Figure 2.**
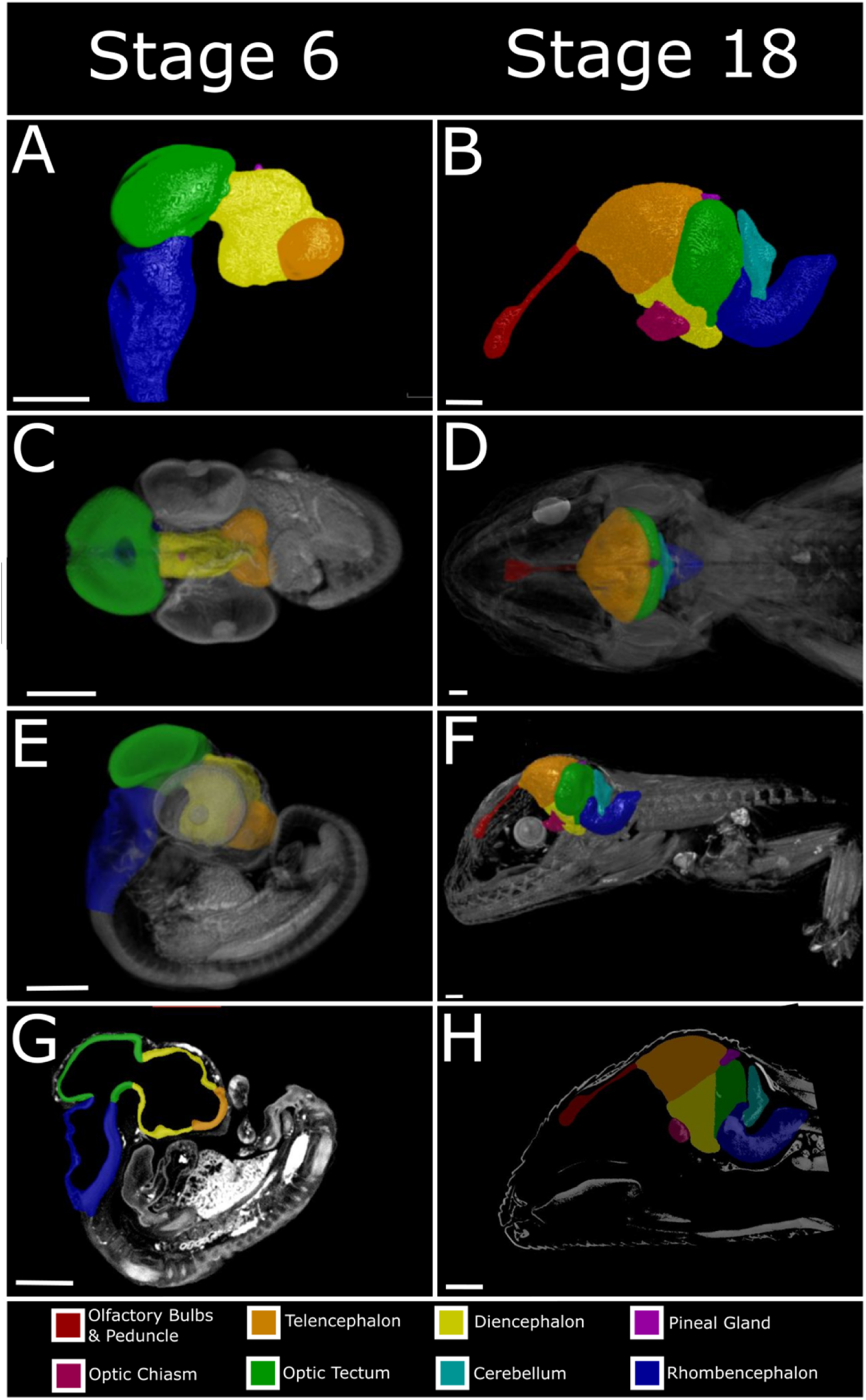
Brains from embryonic stage 6 (left column) and 18 (right column) central bearded dragons (*Pogona vitticeps*), colour coded to highlight the brain regions and associated structures used to characterise brain morphology across stages. Views are as follows: **A-B** lateral, *ex vivo*; **C-D** dorsal, *in situ*, **E-F** lateral, *in situ*, and **G-H** sagittal section *in situ*, highlighting the dramatic expansion of tissue thickness between early and late embryogenesis. Note that the olfactory bulbs and peduncle and the cerebellum are technically part of the telencephalon and rhombencephalon, respectively. Nonetheless, these structures are highly structurally distinct and therefore useful to highlight and discuss for the purpose of characterizing brain development. Scale bars (white, lower left corner) are 1 cm.

**Figure 3.**
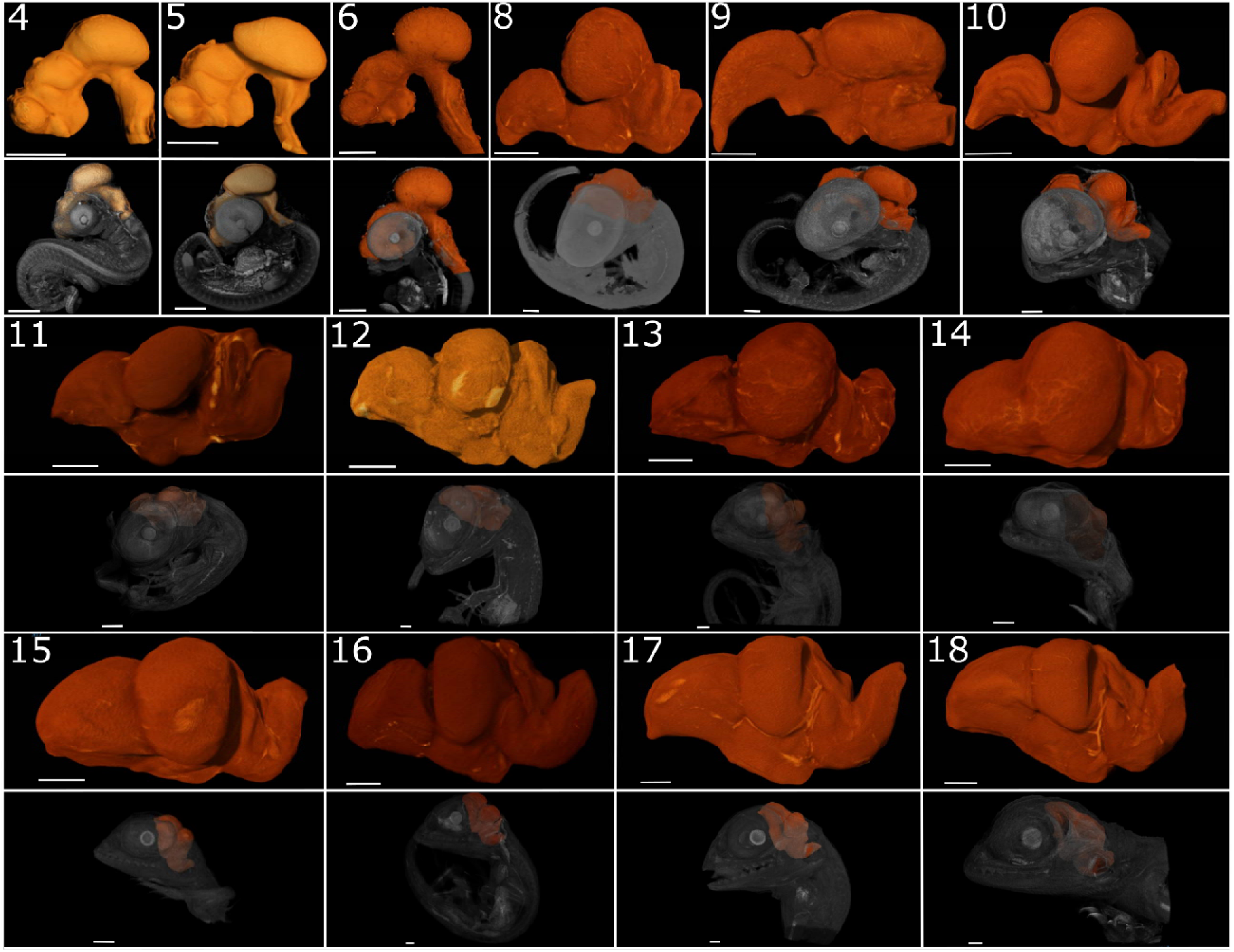
Embryonic development of the central bearded dragon (*Pogona vitticeps*) brain as visualized through diffusible iodine-contrast enhanced µCT (diceCT). The developmental stage is indicated in the top left corner of each brain image and a corresponding image of the brain *in situ* in the embryo is included below. See Figure 2 for representative brain images with the different brain regions used to characterise each stage indicated by colour. Scale bars (white, lower left corner) are 1 cm.

**Table 1:** Neural developmental staging table for *Pogona vitticeps,* based on the Whiteley et al. [2017] embryo staging guide.

| Stage | Description |
| --- | --- |
| 4 | The telencephalon, diencephalon, mesencephalon, and rhombencephalon are all distinguishable. The telencephalon is rounded and laterally compressed. The diencephalon is visible as an elongated, tube-like structure emerging posterior to the telencephalon, with the hypothalamus visible as a ventral extension of the diencephalon. Lateral grooves in the diencephalon differentiate prosomeres 1, 2, and 3. The pineal gland is visible as a bouton on the dorsal midline of the diencephalon. The mesencephalic optic tectum forms a distinct, single ovaline |
|  | lobe approximately four times larger than the telencephalon. The longitudinal fissure is visible along the midline of the telencephalon and the optic tecta. The rhombencephalon appears as a single, tubular structure with a deep neural groove along its dorsal surface. |
| 5 | The telencephalon and the optic tectum are each divided into two lobes due to a deepening of the longitudinal fissure. Both structures are larger, with the telencephalon gaining more prominent lateral extensions. The ventral diencephalon (hypothalamus) has extended ventrally and forms a distinct bulb. The diencephalic prosomeres are distinguishable but have reduced prominence. The optic tecta have begun to fold over the basal mesencephalon. The cerebellum remains fused to the rest of the rhombencephalon but has a distinctive dorsal projection. The neural groove has decreased in prominence. |
| 6 | The telencephalic lobes are larger, with prominent posterolateral extensions. A fissure has formed differentiating the cortex from the subcortex. The ventral diencephalon has extended ventrally and posteriorly towards the ventral surface of the hindbrain. The prosomeres are only subtly distinguishable or no longer discernable along the lateral walls of the diencephalon. The pineal gland is visible as a large bouton on the dorsal surface of the diencephalon, which may be almost as wide as the width of the diencephalon. The optic tectal lobes are larger, extending over the anterior hindbrain and show distinct lateral folding over the ventral mesencephalon. The cerebellum has enlarged and become distinct, with an obvious lateral expansion and flattening, but is still fully connected to the neural tube. The neural groove is absent or almost absent. |
| 8 | The telencephalic lobes extend anteriorly and posteriorly. The anterior-most region of the telencephalon forms a distinct point, which is the developing olfactory bulb. The hypothalamus extends postero-ventrally and can be in contact with the ventral surface of the rhombencephalon. The dorsal surface of the diencephalon is no longer visible, but is covered by the posterior telencephala, pineal gland, and anterior optic tecta. The cerebellum is a distinct tissue extension emerging from the anterior dorsal surface of the rhombencephalon. The neural groove is fully fused. The rhombencephalon, previously straight, has started to |
|  | curve dorsally. |
| **9** | The telencephalon has extended postero-laterally and antero-ventrally. The olfactory bulbs have begun to separate from the rest of the telencephalon, with a fissure forming to mark the boundary between the two structures. The longitudinal fissure along the lateral surface of the telencephalon is deep and pronounced, with the cortex bulging laterally, creating a distinct structure from the subcortex. The diencephalon is completely obscured along the dorsal surface, with the telencephalon and optic tectum approaching or coming in contact lateral to the pineal gland. The dorsal diencephalon (future thalamus) is mostly obscured along the lateral surface. The hypothalamus extends postero-ventrally until it is in contact or almost in contact with the ventral surface of the rhombencephalon. The optic tecta are large and have extended antero-laterally. The cerebellum has formed the beginnings of a distinct tissue layer. The basal rhombencephalon is oriented dorsally and its dorsal surface is pressed against the cerebellum. |
| **10** | The olfactory bulbs are a distinct structure anterior to the telencephalon; the two structures are connected by a well-developed olfactory peduncle. The posterior telencephalon bulges and contacts the anterior optic tectum. The pineal gland is a prominent structure at the midline between the telencephalon and the optic tectum. The dorsal diencephalon is no longer visible along the lateral surface of the brain, being obscured behind the telencephalon and optic tectum, which are in contact. The tuberal hypothalamus has begun to form a distinct structure from the remaining basal diencephalon. The cerebellum forms a distinct, prominent tissue layer pressed against the posterior region of the optic tectum. At the transition from the rhombencephalon to the spinal cord, there is a sharp “knee” bend where the neural tube changes orientation from dorso-ventral to antero-posterior. |
| **11** | The olfactory bulbs have fully migrated to their adult position antero-medial to the eyes and are connected to the telencephalon by the thin olfactory peduncle. The telencephalon has enlarged and the fissure separating the cortex from subcortex is less pronounced. The posterior end of the cortex has begun to fold over the |
|  | subcortex. The optic lobes have increased in volume and expanded laterally, protruding dorsally well beyond the level of the pineal gland and telencephalon. The cerebellum has extended dorsally. |
| **12** | The telencephalon and optic tectum have enlarged but maintained the same overall shape. The fissure between the cortex and subcortex is no longer visible. The cerebellum has extended dorsally, approaching the height of the optic tectum. |
| **13** | The cortex has fully descended ventro-laterally, obscuring the subcortex from a lateral view. The telencephalon and optic tectum have enlarged. The cerebellum has extended dorsally, almost reaching the height of the optic tectum. |
| **14** | The telencephalon and the optic tectum are approximately equal in volume. The optic tectum has extended laterally. The cerebellum has extended dorsally, achieving close to same height as the optic tectum, and has started to cover the postero-dorsal surface of the optic tectum. |
| **15** | The telencephalon is larger than the optic tectum. The cerebellum has begun to thicken at its base. |
| **16** | The cerebellum has thickened substantially, particularly at its base. It now extends posteriorly from the base as well as antero-dorsally. |
| **17** | The cerebellum has reached the height of the optic tectum and lies over the postero-dorsal surface of the optic tectum. The flocculus of the cerebellum is prominent. |
| **18** | The cerebellum extends over the dorsomedial surface of the optic tectum. Along the midline it reaches, or almost reaches, the level of the pineal gland and posterior telencephalon. |

### Brain size variation across incubation temperature

We did not observe an effect of incubation temperature on telencephalon size (Figure 4a; Statistics Supplement) or the size of the rest of the brain (Figure 4b; Statistics Supplement). However, we did observe an effect of genotype on brain size. We found that ZZ genotype embryos had larger telencephala compared to ZW genotype embryos, regardless of incubation temperature (Figure 4c; Statistics Supplement). Although there was no effect of genotype on the size of the rest of the brain, the effect was marginal (Figure 4c; Statistics Supplement). There was no relationship between sexual phenotype and brain size (Supplementary Figure 4). For comparative purposes, Figure 4, panels A and B include vertical dashed lines indicating the developmental stage at which the ovaries and testes begin to differentiate: stages 8 and 9, respectively [Whiteley et al., 2021].

**Figure 4:**
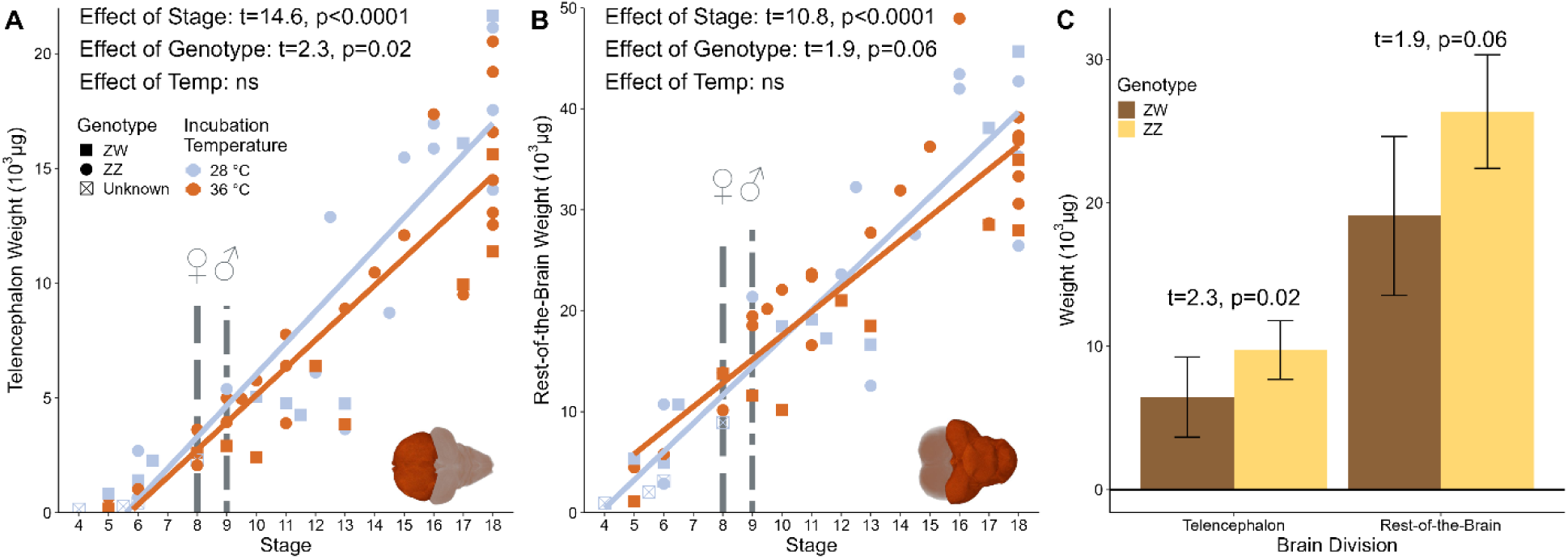
Relationship between developmental stage of embryo, genotype, and brain size across cool (28 °C, blue) and warm (36 °C, orange) incubation temperatures. **A.** Telencephalon mass increases across developmental stages and is greater in ZZ-genotype individuals compared to ZW-genotype individuals. There is no effect of incubation temperature. **B.** The mass of the brain, excluding the telencephalon, increases across developmental stages. There is no effect of temperature or genotype. **A.-B.** Vertical lines indicate gonad differentiation, which begins at stage 8 for ovaries (gray dashed line) and stage 9 for testes (gray double-dashed line) [Whiteley et al., 2017]. Hatching occurs at stage 18. **C.** ZZ genotype embryos have larger brains than ZW genotype embryos, however this effect is only significant in the telencephalon. In the rest of the brain, the effect is marginally non-significant. ns, not significant, i.e. p > 0.05.

Linear models with embryo weight as the predictor revealed that the allometric exponents for the telencephalon and the rest-of-the-brain were similar, and within each other’s 95% confidence intervals, indicating that the telencephalon and the rest-of-the-brain develop at similar rates relative to the overall growth of the embryo (Figure 5; Statistics Supplement). The exponents are less than one, but only marginally, indicating slightly hypoallometric brain growth relative to the rest of the embryo.

**Figure 5.**
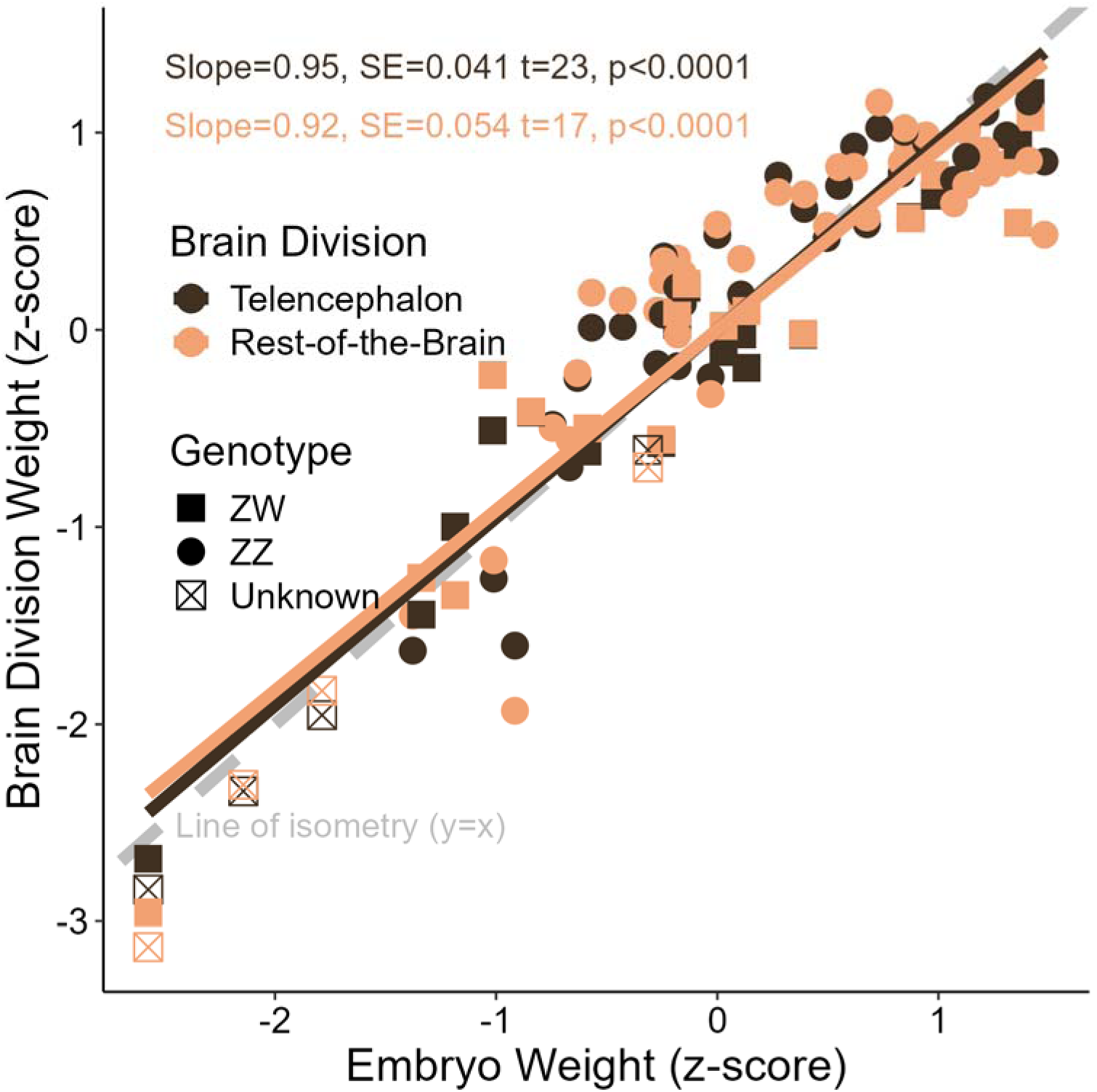
Allometric growth of the central bearded dragon (*Pogona vitticeps*) brain during embryogenesis. There is no difference between the allometric growth rates of the telencephalon and the rest of the brain. Both regions are marginally hypoallometric, increasing in size slightly less rapidly than the embryo overall, as demonstrated by comparing their slopes to the line of isometric growth (grey dashed line). The lines of allometry for the two brain divisions are shallower than the line of isometry.

### Brain shape variation with incubation size

Our linear measurements, transformed using the log-shape ratio and analysed with a permutational ANOVA, reveal that both stage and temperature have significant effects on the shapes of the brains (Figure 6b, Statistics Supplement). There was no significant effect of genotype (Figure 6b; Statistics Supplement) or phenotype (Supplementary Table 1) on brain shape.

**Figure 6.**
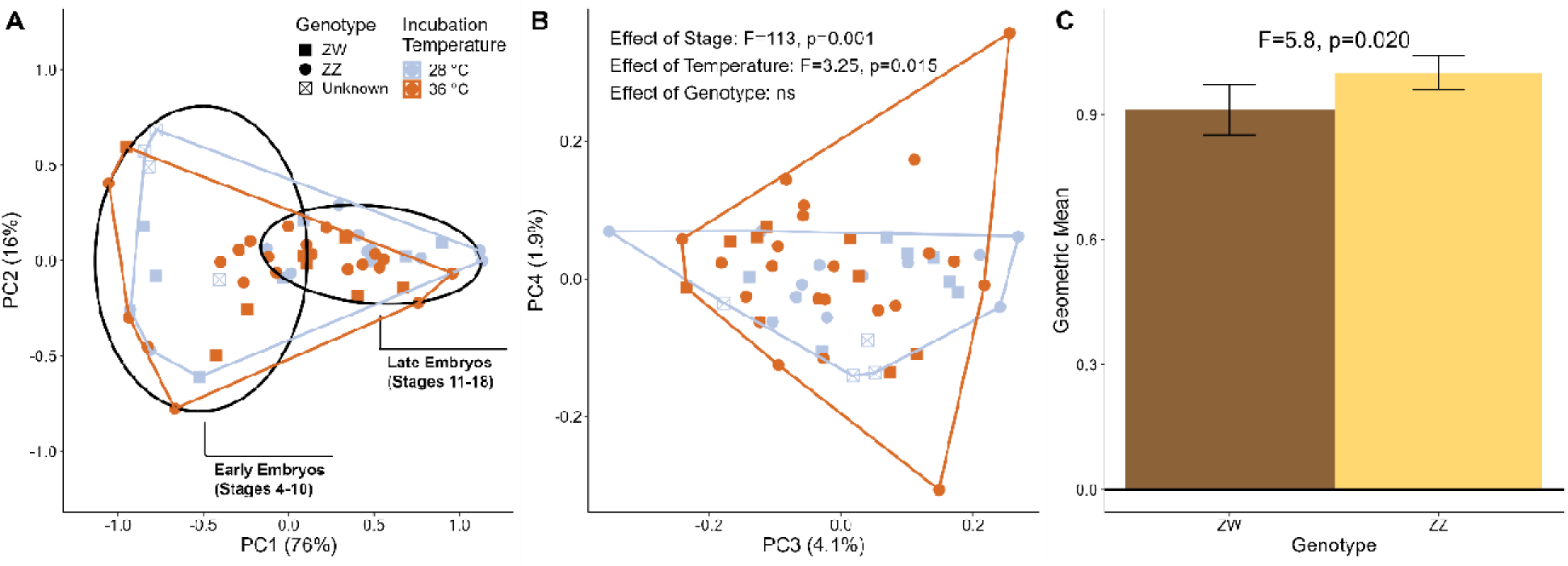
Shape analysis of central bearded dragon (*Pogona vitticeps*) embryonic brains. **A.** The first two components of a principal components analysis of log shape ratios of linear brain measurements reveal that embryo brains change shape as they progress through developmental stages, as reflected along principal component (PC) 1. **B.** The third and forth components of a principal components analysis reveal that warm-incubated embryos occupy a greater volume of morphospace compared to the cool-incubated embryos, particularly along PC4. Inset text shows the results of a permutational ANOVA on the same data, which reveals significant effects of stage and temperature but not genotype. **A.-B.** Replications of these plots with each data point labelled with the stage of the embryo are presented in our Statistics Supplement. **C.** Comparing geometric means reveals a significant effect of genotype. ZZ-genotype embryos have larger brains compared to ZW-genotype embryos. ns = not significant, i.e. p>0.05.

In a plot of the first two components of a principal components analysis of brain shape (Figure 6a), we observe that principal component (PC) 1 is correlated with embryo stage, and a post hoc linear model confirms that this relationship is significant (F=-21.31, p<0.0001; Statistics Supplement). To gain a better understanding of the developmental period during which brains are susceptible to incubation temperature, we divided our brain shape data into two cohorts, early (stages 4–10) and late (stages 11–18) and repeated our permutational ANOVA analysis. We did not find an effect of incubation temperature on the shape of early brains (F=0.37, p=0.36; Supplementary Table 2) but we did find an effect on late brains (F=5.17, p=0.009; Supplementary Table 3). This suggests that brain development is sensitive to elevated incubation temperatures only later in development.

We also found a significant effect of incubation temperature on brain structure (Figure 6b). By plotting the third and fourth components of a principal components analysis, we observe that the warm-incubated embryos occupy a larger volume of morphospace, particularly along PC 4, indicating that these brains show more variation in shape compared to the cool-incubated embryos (Figure 6b). An examination of the principal component analysis loadings (Statistics Supplement) reveals that variation in PC4 is driven primarily by the widths of the telencephalon and optic tectum, the two ovaline structures that emerge early during embryogenesis and remain large and distinct throughout development. Post hoc ANOVAs revealed a significant relationship of incubation temperature with optic tectum width (F=2.2, p=0.032, Supplementary Figure 5) but not telencephalon width (Supplementary Figure 6).

A comparison of geometric means revealed a significant effect of genotype (Figure 6c; Statistics Supplement); the ZZ genotype embryos had larger brains than the ZW genotype embryos, replicating our finding that the telencephalon is larger in ZZ genotype embryos and that there is a trend towards the same for the rest of the brain (Figure 4c). There was no effect of incubation temperature (Statistics Supplement) or phenotype (Supplementary Figure 7) on geometric mean.

## Discussion

We explored how an incubation temperature, warm enough to override genetically determined sexual differentiation, influences embryonic brain development in the central bearded dragon (*Pogona vitticeps)*. We found that the warm incubation temperature affected the shape, but not the size, of the developing brain. We also found that the telencephalon, a region of the forebrain, was larger in embryos with ZZ sex chromosomes compared to ZW sex chromosomes, but this effect could not be teased apart between sexual phenotypes of ZZ males or ZZ sex-reversed females. We discuss the potential implications of these findings for behaviour and cognition in hatchling/adult *P. vitticeps*, as well as broader relevance to how warming climates might impact thermosensitive species.

### Warm incubation temperatures impact body and brain growth in embryonic dragons

We predicted that warm-incubation temperatures would impact brain growth in embryonic dragons due to increased demand for cellular oxygen at higher metabolic rates during development. Since lizards, like all ectotherms, rely on external temperatures to regulate their metabolic activity, warmer incubation temperatures impact the growth rate of embryos [Whiteley et al., 2017], with expected knock-on effects for neurogenesis and other aspects of brain development. In other ectotherms, warmer temperatures have been shown to increase body and brain growth [Andreassen et al., 2022; Gillooly, and McCoy, 2014; Song et al., 2023]. We show that *P. vitticeps* embryos exposed to warmer incubation temperatures (36 °C) hatch in less time than cooler incubated embryos–a mean of 47 days compared to 73 days–with warmer incubated embryos progressing quickly through brain developmental stages (Figure 1b). However, these stimulating effects of heat can reach a threshold, at which point extreme incubation temperatures may have detrimental effects on body and brain growth [Gardner et al., 2011; Beltrán et al., 2024]. Indeed, overall body mass is slightly smaller in warm incubated embryos across all developmental stages in our dataset (Figure 1C), and this trend is consistent with the larger sampling of embryos (n = 234) investigated by Whiteley et al. [2017] (Supplementary Figure 2). This may reflect a broader trend of climate warming and smaller body sizes among ectotherms [Ohlberger, 2013].

The impact of heat stress on brain development has been investigated in other lizards. Extreme incubation temperature (33–36 °C) in brown anoles (*Anolis sagrei*) impacts survivorship and induce craniofacial abnormalities that encroach on the internal cranial space, causing cell death and stunting telencephalon growth [Sanger et al., 2021]. Similarly, a higher-than-normal incubation temperature (24 °C versus 18 °C) in alpine-dwelling Eastern three-lined skinks (*.Acritoscincus duperreyi*) reduces telencephalon size [Amiel et al., 2017]. Functionally, elevated incubation temperatures alter the cognition and behaviour of lizards [Beltrán et al., 2024; Dayananda, and Webb, 2017; Siviter et al., 2017a]. In our dataset, however, comparable warm incubation temperatures (36 °C) had more subtle effects on neural development that did not impact brain size in embryonic *P. vitticeps* dragons (Figure 3). This may be due to acclimatory processes associated with local adaptation because *P. vitticeps* is an arid specialist that are often exposed to high nesting temperatures across their central Australian desert range [Castelli et al., 2021]. Although incubation temperatures above 32 °C can induce sex-reversal in *P. vitticeps*, this treatment is not extreme enough to induce any pathological behavioural or somatic morphological effects [Whiteley et al., 2017], so it is not likely to be extreme enough to induce the broad, substantial changes to brain development as has been seen in tropical- and alpine-adapted lizards such as *Anolis* and *Acritoscincus*, respectively.

In fact, we expect that temperature-induced variation in brain development may be more widespread than we have found but that the effects are too subtle to detect using gross anatomical measurements of external brain morphology. Indeed, evolutionary changes in brain anatomy can be specific to individual nuclei [Hoops et al., 2022; Hoops et al., 2017a] and/or occur in a mosaic fashion such that compensatory changes across different brain regions effectively cancel out any effects that would be detectable at the gross level [Hoops et al., 2017b]. Our linear shape analysis revealed a difference in the shapes of brains incubated different temperatures (Figure 6b, Statistics Supplement), however this effect did not translate into detectable volumetric differences (Figure 4c). It is likely that temperature-induced changes are being induced at the level of individual nuclei and fibre tracts throughout the brain, and our finding of a wider optic tectum is just the first example. These specific, localized differences can result in pervasive yet subtle changes in brain function. Furthermore, the mechanisms underlying these changes may be either adaptive, reflecting protective neurodevelopmental responses to elevated temperatures, or pathological, arising from temperature-induced dysregulation of developmental processes. Detecting and characterizing these effects will be challenging and require comprehensive behavioural testing to generate hypotheses and multimodal neuroanalytical approaches to detect differences at anatomical, cellular, and molecular levels.

### Larger telencephala in embryos with ZZ genotypes indicates greater influence of genetically driven sex determination on brain growth

We found significantly larger telencephala, one of the two primary divisions of the forebrain, in individuals with the ZZ genotype (ZZf, ZZm) compared to the ZW genotype (ZWf), while the rest-of-the brain was no different in size across the two genotypes and three sexual phenotypes (Figure 4, Supplementary Figure 4). These results contrast previous studies that show distinct developmental cascades within the gonads between genetically-driven and thermally-driven females in *P. vitticeps* [Whiteley et al., 2021]. Squamates have an unusually long duration of gonad differentiation during development, with prolonged stages of temporary pseudohermaphroditism (stage 10–17) and late hemipene regression (stage 17) in *P. vitticeps* [Whiteley et al., 2018; Whiteley et al., 2017]. In contrast, differentiation of major brain regions progressed quickly in early embryonic stages (stages 4–10; Figure 3; Figure 4) before shifting into steady allometric growth across the brain regions up until hatching (stages 11–18; Figure 3; Figure 4). Qualitative examination of brain mass in our dataset suggests that the timing of growth for the telencephalon and the rest-of-the-brain may be delayed until the onset of gonad differentiation (Figure 4, A-B vertical lines). Note the unusual clustering of points below the regression lines at stages 10–13 in Figure 4 A-B. However, our limited sample size at these stages across cool and warm incubation temperatures precluded statistical testing, so this result should be considered preliminary. Our results raise questions about the developmental timing of brain differentiation with potential implications for understanding the role of the crosstalk between the brain and gonads via the hypothalamus-pituitary axis.

Post hatching, several phenotypic differences have been observed for sex-reversed female (ZZf) compared to male (ZZm) *P. vitticeps*. Captive studies have shown that ZZf individuals have higher fecundity [Holleley et al., 2015] and a unique phenotype “male-like” longer relative tail lengths and higher preferred body temperatures, “female-like” larger body sizes and bite forces, and some unique traits, including personalities and metabolic rates distinct from concordant females and males [Li et al., 2016; Wild et al., 2023; Jones et al., 2020]. Furthermore, a recent study on wild, free roaming bearded dragon populations found that individual movement patterns, fecundity and body condition were no different between sex reversed and concordant females [Wild et al., 2022]. Significantly larger telencephala in ZZm and ZZf might influence behavioural traits such as boldness, but behavioural studies to directly test cognition in ZZ sexes compared to typical ZW females are needed to investigate whether embryonic differences in telencephalon sizes are linked with behavioural differences into adulthood. Post-hatch characterization of neuroanatomy across genotypes is also needed to identify the adult neural implications of larger telencephala during embryonic development.

### Do differences in brain size reflect functional changes in neurons and/or behaviour?

A key assumption in comparative neurobiology is that brain size reflects greater neuronal investment, such as greater neuronal density or more extensive connectivity, which in turn influences behavioural traits under selection, including social cognition and sensory processing [Jerison, 1973]. This relationship between telencephalon size and performance is somewhat reflected in alpine skinks, A*. duperreyi,* whereby hatchlings and juveniles incubated at warmer temperatures were faster learners and sometimes performed better in cognitive tasks; linked with differences in volume of the telencephala and size and density of neurons within this brain region [Clark et al., 2013; Amiel et al., 2014; Amiel, and Shine, 2012; Amiel et al., 2017]. Interestingly, social learning in adult *P. vitticeps* was impacted by incubation temperatures in a previous study with cool-incubated dragons (27 ± 3°C) learning social tasks faster than warm-incubated dragons (30 ± 3 °C) [Siviter et al., 2017b]. These studies were all conducted prior to the discovery of temperature-induced sex-reversal in these species, so neither genotypic nor phenotypic sex were accounted for as potential confounding factors. New studies are needed that account for sex-reversal effects when examining these questions.

Although brain size, neuron density, connectivity, and brain function (i.e. cognition and behaviour) are linked, the relationship between them is not absolute and examining only one of these characters only provides an incomplete picture of neural changes. For example, velvet geckos (*Amalosia leseurii*) show temperature-induced alterations in neural development, in this case smaller telencephala, but they showed no differences in neuronal density or learning ability as adults [Beltrán et al., 2024; Abayarathna, and Webb, 2020]. In another example, *Anolis* lizards show between-species variation in neuron density that is not reflected volumetrically (behaviour was not tested) [Leal, and Powell, 2012; Storks et al., 2020; Storks et al., 2024]. This highlights how subtle the functional effects of even gross morphological changes to brain anatomy can be. The telencephalon has a myriad of diverse roles and pinpointing functional implications, particularly sub-pathological ones, can be challenging. Nonetheless, it is likely that any temperature-induced alteration in brain morphology has some functional implications; the challenge lies in determining just what those functional implications may be, and what effects they may have on individual fitness.

### Evolutionary consequences of altered brain development in dragons

Our results suggest that an abnormally warm incubation temperature does alter brain development in *P. vitticeps* although the effect is subtle and was detected only in our shape analysis. Due to the specificity of this effect, it is likely to affect only certain brain regions, such as the optic tectum, i.e. alter brain development in a mosaic fashion [Barton, and Harvey, 2000]. The function(s) of these regions will dictate the nature, pervasiveness, and fitness consequences of altered brain development under warming climate scenarios, and ultimately determine, in part, how climate change can alter the evolutionary trajectory of ectothermic species. In the case of the optic tectum, effects are most likely to be in visual processing and vision-mediated behaviour. Identifying more affected regions, by quantifying volumetric changes in specific brain regions, will be an important step forward in our understanding of how the ectothermic brain responds to thermal challenges.

Our study did not detect any neurodevelopmental effects specific to the sex-reversed ZZf phenotype. However, as we note above, this phenotype is associated with a unique behavioural profile that is distinct from the two concordant sexes (ZWf and ZZm) [Li et al., 2016]. Some of these traits are linked to reproductive behaviour, such as increased fecundity, which indicate possible alterations in the endocrine system. We expect that there are neurodevelopmental effects of sex-reversal, likely in regions involved in endocrine system regulation and sex-specific behaviour, and we hypothesize that these effects are more subtle than we could detect here and/or are initiated during post-hatch neural development. Therefore, this sex-reversal mechanism may represent a valuable model for teasing apart the complex interplay of genetic, environmental, and endocrine influences on the maturation and sexual differentiation of the brain. Brain regions that are involved in reproductive function and sex-specific behaviours, such as the medial preoptic area and ventromedial hypothalamus [Numan, 2014], are among the most likely potential targets for localized neural differences between ZWf, ZZm, and ZZf phenotypes.

### A detailed neuroanatomy staging guide for dragons

The central bearded dragon is emerging as a model organism with applications across anatomy and physiology [Fenk et al., 2024]. It is particularly important for developmental studies and, as such, the characterization and standardization of its developmental trajectory is a critical step in moving the field forward. An overall characterization of bearded dragon embryonic development based on limb and genital morphology has been published [Whiteley et al., 2017], as has a description of their cranial development [Ollonen et al., 2018].

In this study, we have generated 57 3D images of *P. vitticeps* embryos, spanning the majority of their post-oviposition, pre-hatch development, resulting in an information-rich anatomical dataset of the entire embryo for early developmental stages and the anterior portion of the embryo for later developmental stages (Figure 3). We used iodine-based contrast-enhanced soft tissue µCT [Collin et al., 2026] to acquire a high-resolution, detailed dataset of embryonic development. Our results represent, to our knowledge, the most complete and detailed 3D imaging dataset of the development of any animal.

Using these data, we present an anatomical guide for the brain development in *P. vitticeps*. Broadly, we differentiate brain development into two phases. During early brain development there are substantial changes to the shape of the brain between stages, while during late brain development changes are less substantial and more characterised by changes in size rather than shape. Early embryonic brain development consists of stages 4–10 and is most readily characterised by the dorsal diencephalon (developing thalamus) being visible from the lateral view, and a relatively straight rhombencephalon at its boundary with the spinal cord (Table 1; Figure 3). Late embryonic brain development consists of stages 11–18 and is most readily characterised by the telencephalon and optic tectum coming in contact, obscuring the dorsal diencephalon from lateral view, and the rhombencephalon forming a sharp “knee” bend where it meets the spinal cord (Table 1; Figure 3). Notably, we found that the effect of incubation temperature on brain shape occurred during late embryonic development, suggesting that the brains are at least somewhat protected from the neural effects of elevated incubation temperature during the first stages of their development, when the most dramatic anatomical changes are occurring.

In *Pogona*, as in all vertebrates, the brain consists of tissue surrounding the cerebral aqueduct, the adult form of the central lumen of the embryonic neural tube, and its associated ventricles [Foss et al., 2022]. There are four large brain divisions that grow to become substantially separate structures to this: the olfactory bulbs, the telencephalon, the optic tectum, and the cerebellum. Of these four structures, two develop during early brain development: the telencephalon and the optic tectum. These structures are already clearly distinguishable at stage 4, the earliest stage we examined (Table 1; Figure 3). The other two structures, the olfactory bulbs and the cerebellum, do not appear as prominent, independent structures until late embryonic brain development (Table 1; Figure 3). The relative sizes of these structures in adulthood do not support the well-known “late equals large” hypothesis [Finlay et al., 2001; Weisbecker, 2009], because in *Pogona* earlier-appearing structures become much larger than the later-appearing structures. Whether this is indicative of a more heterochronic pattern of neurogenesis in *Pogona* compared to mammalian embryonic development, where the late equals large hypothesis originates, needs to be investigated.

Staging guides for embryonic development are now available for several lizard species [e.g. see summary table in Ollonen et al., 2018]. These guides are traditionally based on external photographs of embryos and frequently include brief descriptions of brain morphology to the extent it is visible, as embryos are transparent for a substantial portion of their development [e.g. Lin et al., 2021; Py Daniel et al., 2017; Wen, and Griffith, 2026]. Although these descriptions are useful, the development of most of the brain cannot be determined because it is obscured behind other developing structures, principally the eyes, and later by the skin. Therefore, the descriptions of brain development in these staging guides are necessarily incomplete.

Our study reinforces the utility of buffered iodine contrast-enhanced bioimaging for generating high resolution, detailed anatomical 3D images of interior soft-tissue structures [Collin et al., 2026; Metscher, 2009b; Lanzetti, and Ekdale, 2021]. Using these images, we have been able to visualize and describe the developing brain in fine detail, noting even relatively subtle anatomical changes between developmental stages (Table 1, Figure 3, 4). To our knowledge, only one other publication has used contrast-enhanced soft-tissue µCT scanning to create an embryonic staging guide of a lizard [Griffing et al., 2019]. As in our *Pogona* guide, they describe brain development in substantial detail across stages. As a result, we can observe that the brain of the mourning gecko (*Lepidodactylus lugubris*) develops largely along the same trajectory as *Pogona*. There are, however, relatively minor differences. For example, the olfactory bulbs appear to develop slightly earlier, and the cerebellum slightly later, in *Lepidodactylus*. Also, we note that Griffing et al. use a stage numbering that commences at fertilization, such that oviposition, our stage 1, is designated by them as stage 27. Stage 30 in *Lepidodactylus* shows about the same level of brain development as stage 4 in *Pogona*, and the final stage before hatching is stage 43 in *Lepidodactylus* and stage 18 in *Pogona*. The transition between early and late brain development, which occurs around stage 10 in *Pogona*, appears to occur around stage 35 in *Lepidodactylus.* The availability of these two 3D-imaging-based stage series represents the first opportunity for a detailed comparative analysis of lizard embryonic development, potentially a rich new avenue of evolutionary developmental research.

### Conclusions

Thermosensitive species like *P. vitticeps* offer a unique opportunity to investigate the interacting effects of heat and sex on brain development in a warming world. Our results indicate that embryonic brain development in dragons may be somewhat buffered to the effects of warmer incubation temperatures. In contrast to previous studies in *P. vitticeps*, the ZZ genotype has a greater impact than thermal conditions with significantly larger telencephala in ZZm and ZZf than concordant ZWf phenotypes. Future studies should aim to use histological techniques to determine the neuronal density and developmental mechanisms that may be driving differences in telencephalon sizes across sexes.

## Acknowledgements

We thank the Melbourne TrACEES (Trace Analysis for Chemical, Earth and Environmental Sciences) Platform for access to the µCT scanner and Jay Black for technical support. We thank Australian Nuclear Science and Technology Organisation Australian Synchrotron for the beamtime granted toward this project and Anton Maksimenko for technical support. Thanks to Alice Clement, Caroline Kerr, Lucille Chapuis, Myoung Hoon Ha and Hope Robins for assistance at the Synchrotron. Thanks to Mike S. Y. Lee, Clare Holleley and Arthur Georges for early discussions on the project design. Thanks to Emma Sherratt for advice on analyzing linear shape data.

## Statement of Ethics

This research used preserved specimens from a previous study so did not require animal ethics.

## Author Contributions

J.M.C-R and S.P.C conceived of the study, S.W. provided the specimens and background data, A.D.C. and J.M.C-R prepared and scanned specimens, A.D.C. segmented the 3D images and collected the data, D.H. prepared the figures and the staging guide, D.H., J.M.C-R. and R.P. analysed the data, J.M.C-R, A.D.C and D.H. wrote the manuscript with input from all co-authors.

## Data Availability Statement

Tomographic stacks for all scans and all data are available via FigShare (https://doi.org/10.25909/c.8789142).

## Supplementary Materials

**Supplementary Material 1: Statistics Supplement:** Full statistical outputs for all the analyses presented in the manuscript.

**Supplementary Material 2: Supplementary Figures & Tables:** Supplementary analyses to completement the principal findings that we present in the manuscript.

