## Supplementary Materials 1 for "Impact of heat and sex on embryonic brain growth in dragons with temperature induced sex-reversal (Pogona vitticeps, Lacertilia: Agamidae)"

Statistics Supplement

Daniel Hoops

This document provides detailed statistical outputs for all the analyses presented in the manuscript “Impact of heat and sex on embryonic brain growth in dragons with temperature induced sex-reversal (Pogona vitticeps, Lacertilia: Agamidae)” by Daniel Hoops, Annalise D. Ceddia, Richard Peters, Sarah Whiteley, Shaun P. Collin, and Jenna M. Crowe-Riddell. Each panel listed below corresponds to a figure panel in the main manuscript.

### Figure 1, panel B

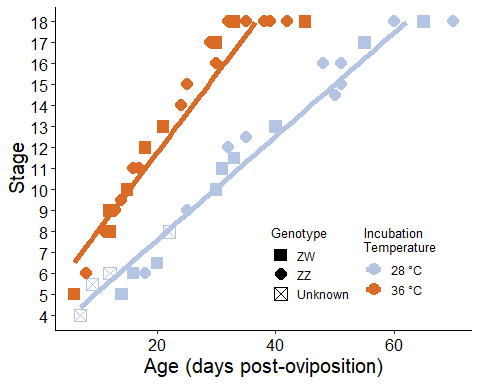

This panel illustrates ages at which embryos reach each developmental stage. Embryos incubated at a warmer temperature progress through developmental stages faster than embryos incubated at a normal (relatively cool) inclubation temperature.

### Figure 1, panel C

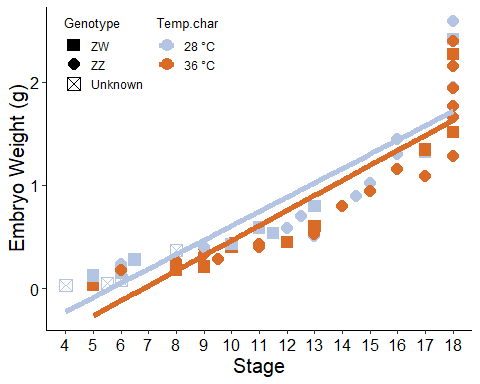

This panel illustrates the difference in the weights of embryos incubated at two different incubation temperatures.

Summary Statistics:

| Temp.char | Genotype | N | Embryo.weight | sd | se | ci |
| --- | --- | --- | --- | --- | --- | --- |
| 28 °C | ZW | 9 | 0.74 | 0.73 | 0.24 | 0.56 |
| 28 °C | ZZ | 13 | 1.04 | 0.73 | 0.20 | 0.44 |
| 28 °C | Unknown | 4 | 0.13 | 0.16 | 0.08 | 0.25 |
| 36 °C | ZW | 9 | 0.78 | 0.76 | 0.25 | 0.58 |
| 36 °C | ZZ | 22 | 0.87 | 0.70 | 0.15 | 0.31 |

Significant differences in the weight of the embryos between temperature treatments and stages were detected using a linear model. We calculated a linear model with temperature, genotype, and stage as fixed effects, and embryo weight as the response variable.

```
##
## Call:
## lm(formula = Emb.w.lncs ~ Stage.cs + Genotype + Incubation.temperature,
##     data = data[data$Genotype != "Unknown", ])
##
## Residuals:
##      Min       1Q   Median       3Q      Max
## -1.10702 -0.10926 -0.00932  0.14052  0.38831
##
## Coefficients:
##                         Estimate Std. Error t value Pr(>|t|)
## (Intercept)             0.794337   0.270034   2.942  0.00497 **
## Stage.cs                0.877278   0.034425  25.484  < 2e-16 ***
## GenotypeZZ              0.111516   0.069556   1.603  0.11531
## Incubation.temperature -0.025350   0.008257  -3.070  0.00349 **
## ---
## Signif. codes:  0 '***' 0.001 '**' 0.01 '*' 0.05 '.' 0.1 ' ' 1
##
## Residual standard error: 0.2351 on 49 degrees of freedom
## Multiple R-squared:  0.9333, Adjusted R-squared:  0.9292
## F-statistic: 228.5 on 3 and 49 DF,  p-value: < 2.2e-16
```

### Figure 4, panel A

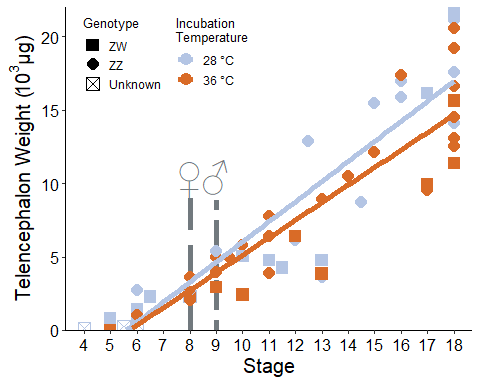

This panel illustrates increasing telencephalon weight during embryogenesis.

Summary Statistics:

| Genotype | Temp.char | N | Tel.weight.ug | sd | se | ci |
| --- | --- | --- | --- | --- | --- | --- |
| ZW | 28 °C | 9 | 6788.10 | 7161.90 | 2387.30 | 5505.12 |
| ZW | 36 °C | 9 | 6140.03 | 5117.44 | 1705.81 | 3933.61 |
| ZZ | 28 °C | 13 | 10858.87 | 6633.93 | 1839.92 | 4008.84 |
| ZZ | 36 °C | 22 | 9082.89 | 5972.13 | 1273.26 | 2647.90 |
| Unknown | 28 °C | 4 | 787.36 | 1015.35 | 507.68 | 1615.65 |

Significant differences in telencephalon weight were detected with respect to genotype and stage but not incubation temperature. We calculated a linear mode with stage, genotype, and incubation temperature as fixed effects and telencephalon weight as the response variable.

```
##
## Call:
## lm(formula = Tel.w.lncs ~ Stage.cs + Genotype + Incubation.temperature,
##     data = data[data$Genotype != "Unknown", ])
##
## Residuals:
##     Min      1Q  Median      3Q     Max
## -1.3408 -0.1838  0.0471  0.2574  0.4700
##
## Coefficients:
##                        Estimate Std. Error t value Pr(>|t|)
## (Intercept)             0.46509    0.41459   1.122   0.2674
## Stage.cs                0.77291    0.05285  14.624   <2e-16 ***
## GenotypeZZ              0.25044    0.10679   2.345   0.0231 *
## Incubation.temperature -0.01722    0.01268  -1.359   0.1805
## ---
## Signif. codes:  0 '***' 0.001 '**' 0.01 '*' 0.05 '.' 0.1 ' ' 1
##
## Residual standard error: 0.361 on 49 degrees of freedom
## Multiple R-squared:  0.828,  Adjusted R-squared:  0.8175
## F-statistic: 78.64 on 3 and 49 DF,  p-value: < 2.2e-16
```

### Figure 4, panel B

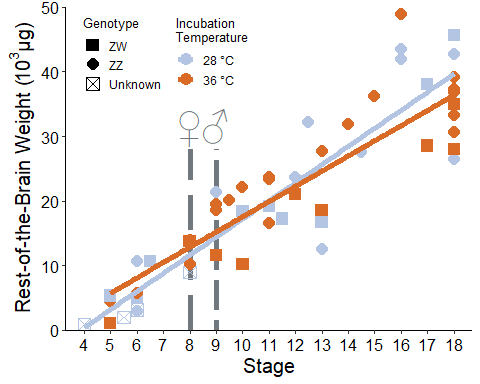

This panel illustrates increasing rest-of-the-brain weight during embryogenesis.

Summary Statistics:

| Genotype | Temp.char | N | Rest.weight.ug | sd | se | ci |
| --- | --- | --- | --- | --- | --- | --- |
| ZW | 28 °C | 9 | 19583.85 | 13855.36 | 4618.45 | 10650.17 |
| ZW | 36 °C | 9 | 18616.92 | 10653.52 | 3551.17 | 8189.02 |
| ZZ | 28 °C | 13 | 27467.55 | 12973.40 | 3598.17 | 7839.75 |
| ZZ | 36 °C | 22 | 25726.23 | 11570.49 | 2466.84 | 5130.07 |
| Unknown | 28 °C | 4 | 3765.86 | 3557.83 | 1778.91 | 5661.30 |

Significant differences in the weight of the rest of the brain were detected with respect to stage but not genotype or incubation temperature. We calculated a linear model with stage, genotype, and incubation temperature as fixed effects and telencephalon weight as the response variable.

```
##
## Call:
## lm(formula = Rest.w.lncs ~ Stage.cs + Genotype + Incubation.temperature,
##     data = data[data$Genotype != "Unknown", ])
##
## Residuals:
##     Min      1Q  Median      3Q     Max
## -1.7512 -0.2096  0.1548  0.3442  0.5181
##
## Coefficients:
##                         Estimate Std. Error t value Pr(>|t|)
## (Intercept)             0.171879   0.514835   0.334   0.7399
## Stage.cs                0.707634   0.065634  10.782 1.56e-14 ***
## GenotypeZZ              0.254885   0.132612   1.922   0.0604 .
## Incubation.temperature -0.008011   0.015743  -0.509   0.6131
## ---
## Signif. codes:  0 '***' 0.001 '**' 0.01 '*' 0.05 '.' 0.1 ' ' 1
##
## Residual standard error: 0.4483 on 49 degrees of freedom
## Multiple R-squared:  0.7253, Adjusted R-squared:  0.7085
## F-statistic: 43.12 on 3 and 49 DF,  p-value: 8.703e-14
```

### Figure 4, panel c

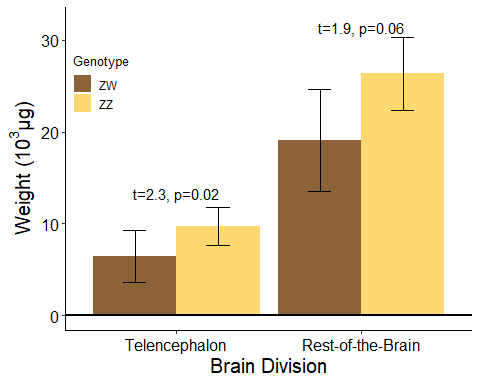

This graph illustrates the differences in weight between genotypes. These results are from the same statistical analyses as Figures 4a and 4b.

### Figure 5

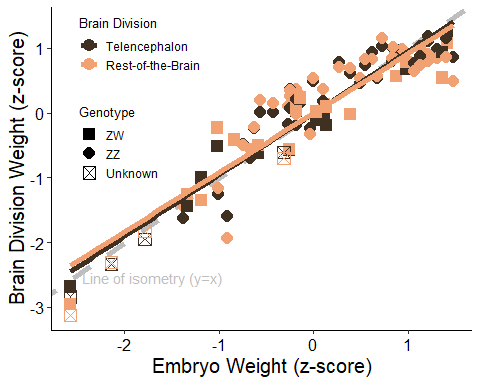

This panel illustrates the allometric relationships between embryo weight and the weights of the two brain divisions.

Linear models reveal that both brain divisions are slightly hypoallometric, i.e. their allometric slopes are slightly less than 1, and the confidence intervals do not overlap with 1.

```
## [1] "Telencephalon:"
##
## Call:
## lm(formula = Tel.w.lncs ~ Emb.w.lncs, data = data)
##
## Residuals:
##      Min       1Q   Median       3Q      Max
## -0.73024 -0.25581 -0.04439  0.23931  0.60259
##
## Coefficients:
##              Estimate Std. Error t value Pr(>|t|)
## (Intercept) 7.590e-16  4.098e-02    0.00        1
## Emb.w.lncs  9.518e-01  4.135e-02   23.02   <2e-16 ***
## ---
## Signif. codes:  0 '***' 0.001 '**' 0.01 '*' 0.05 '.' 0.1 ' ' 1
##
## Residual standard error: 0.3094 on 55 degrees of freedom
## Multiple R-squared:  0.906,  Adjusted R-squared:  0.9043
## F-statistic:   530 on 1 and 55 DF,  p-value: < 2.2e-16
## [1] "Rest-of-the-Brain:"
##
## Call:
## lm(formula = Rest.w.lncs ~ Emb.w.lncs, data = data)
##
## Residuals:
##      Min       1Q   Median       3Q      Max
## -1.09423 -0.29634  0.02927  0.32765  0.70830
##
## Coefficients:
##              Estimate Std. Error t value Pr(>|t|)
## (Intercept) 5.782e-16  5.389e-02    0.00        1
## Emb.w.lncs  9.151e-01  5.437e-02   16.83   <2e-16 ***
## ---
## Signif. codes:  0 '***' 0.001 '**' 0.01 '*' 0.05 '.' 0.1 ' ' 1
##
## Residual standard error: 0.4068 on 55 degrees of freedom
## Multiple R-squared:  0.8374, Adjusted R-squared:  0.8345
## F-statistic: 283.3 on 1 and 55 DF,  p-value: < 2.2e-16
```

### Figure 6, panels A & B

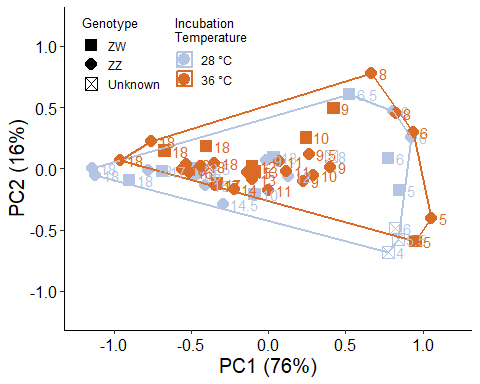

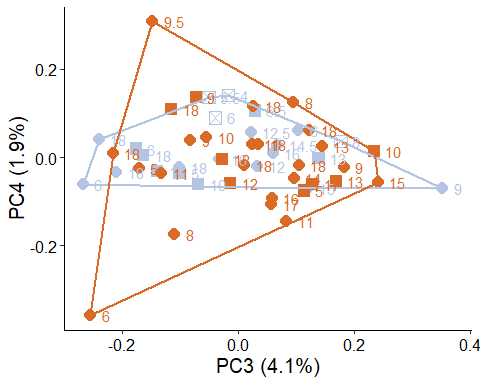

These panels illustrate the morphospace occupied by the brains.

The linear measurements used are: LB: brain length LET_L: left telencephalon length LET_R: right telencephalon length MWT: telencephalon width LEM_L: left mesencephalon length LEM_R: right mesencephalon length MWM: mesencephalon width MDD: diencephalon depth

Variance explained by components:

|  | PC1 | PC2 | PC3 | PC4 | PC5 | PC6 | PC7 | PC8 |
| --- | --- | --- | --- | --- | --- | --- | --- | --- |
| Standard deviation | 0.60 | 0.27 | 0.14 | 0.10 | 0.07 | 0.05 | 0.03 | 0 |
| Proportion of Variance | 0.76 | 0.16 | 0.04 | 0.02 | 0.01 | 0.01 | 0.00 | 0 |
| Cumulative Proportion | 0.76 | 0.92 | 0.96 | 0.98 | 0.99 | 1.00 | 1.00 | 1 |

PC loadings:

|  | PC1 | PC2 | PC3 | PC4 | PC5 | PC6 | PC7 | PC8 |
| --- | --- | --- | --- | --- | --- | --- | --- | --- |
| LB | -0.05 | 0.06 | 0.08 | 0.18 | 0.90 | -0.15 | 0.08 | -0.35 |
| LET_L | -0.14 | 0.53 | -0.25 | 0.03 | -0.07 | 0.17 | -0.69 | -0.35 |
| LET_R | -0.14 | 0.47 | -0.31 | 0.09 | -0.25 | -0.10 | 0.68 | -0.35 |
| MWT | -0.45 | -0.23 | 0.54 | 0.49 | -0.27 | 0.15 | -0.02 | -0.35 |
| LEM_L | 0.57 | -0.13 | 0.05 | 0.21 | -0.24 | -0.64 | -0.17 | -0.35 |
| LEM_R | 0.59 | -0.10 | 0.05 | -0.03 | 0.00 | 0.70 | 0.13 | -0.35 |
| MWM | -0.12 | 0.05 | 0.44 | -0.80 | -0.05 | -0.14 | 0.03 | -0.35 |
| MDD | -0.28 | -0.65 | -0.59 | -0.17 | -0.02 | 0.01 | -0.05 | -0.35 |

We used a permutational ANOVA to analyse the shape measurement with respect to stage, incubation temperature, and genotype. Stage and incubation temperature, but not genotype, where significant.

```
##
## Analysis of Variance, using Residual Randomization
## Permutation procedure: Randomization of null model residuals
## Number of permutations: 1000
## Estimation method: Ordinary Least Squares
## Sums of Squares and Cross-products: Type I
## Effect sizes (Z) based on F distributions
##
##           Df      SS      MS     Rsq        F       Z Pr(>F)
## Stage      1 14.8177 14.8177 0.69995 112.8053  5.3617  0.001 ***
## Genotype   1  0.0136  0.0136 0.00064   0.1032 -2.3861  0.988
## Temp       1  0.4272  0.4272 0.02018   3.2522  2.0126  0.024 *
## Residuals 45  5.9110  0.1314 0.27922
## Total     48 21.1695
## ---
## Signif. codes:  0 '***' 0.001 '**' 0.01 '*' 0.05 '.' 0.1 ' ' 1
##
## Call: procD.lm(f1 = lsr ~ Stage + Genotype +
## Temp, data = analyze_lsr_gen)
```

This post hoc linear model demonstrates that there is an effect of stage on the first principal component of a principal components analysis of brain shape.

```
##
## Call:
## lm(formula = PC1 ~ Stage, data = plotPCs)
##
## Residuals:
##     Min      1Q  Median      3Q     Max
## -0.4344 -0.0945  0.0049  0.1413  0.3645
##
## Coefficients:
##              Estimate Std. Error t value Pr(>|t|)
## (Intercept)  1.469514   0.073885   19.89   <2e-16 ***
## Stage       -0.121221   0.005688  -21.31   <2e-16 ***
## ---
## Signif. codes:  0 '***' 0.001 '**' 0.01 '*' 0.05 '.' 0.1 ' ' 1
##
## Residual standard error: 0.1933 on 51 degrees of freedom
## Multiple R-squared:  0.8991, Adjusted R-squared:  0.8971
## F-statistic: 454.2 on 1 and 51 DF,  p-value: < 2.2e-16
```

### Figure 6, panel c

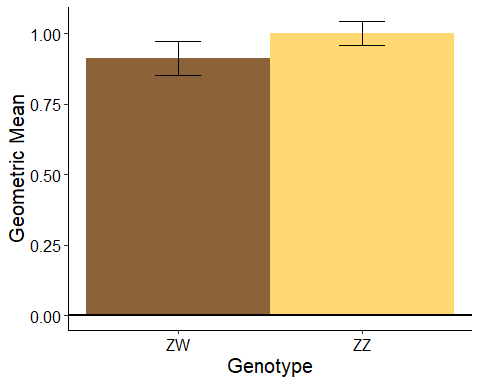

This panel illustrates the brain geometric mean (a measure of brain size) with respect to genotype.

Summary Statistics:

| Genotype | Temp.char | N | geomean | sd | se | ci |
| --- | --- | --- | --- | --- | --- | --- |
| ZW | 28 °C | 7 | 0.90 | 0.22 | 0.08 | 0.21 |
| ZW | 36 °C | 8 | 0.87 | 0.31 | 0.11 | 0.26 |
| ZZ | 28 °C | 12 | 1.02 | 0.18 | 0.05 | 0.11 |
| ZZ | 36 °C | 22 | 1.00 | 0.15 | 0.03 | 0.07 |
| Unknown | 28 °C | 4 | 0.33 | 0.31 | 0.15 | 0.49 |

An ANOVA reveals significant effects of stage and genotype, but not incubation temperature, on brain geometric mean.

```
## Anova Table (Type II tests)
##
## Response: geomean
##            Sum Sq Df F value    Pr(>F)
## Stage     1.19822  1 89.8750 2.694e-12 ***
## Genotype  0.07768  1  5.8269   0.01991 *
## Temp.char 0.01190  1  0.8924   0.34986
## Residuals 0.59994 45
## ---
## Signif. codes:  0 '***' 0.001 '**' 0.01 '*' 0.05 '.' 0.1 ' ' 1
```
