## Supplementary Materials 2 for "Impact of heat and sex on embryonic brain growth in dragons with temperature induced sex-reversal (Pogona vitticeps, Lacertilia: Agamidae)"

Supplementary Figures & Tables

Daniel Hoops

This document provides supplementary statistical analyses that complement the analyses presented in the manuscript “Impact of heat and sex on embryonic brain growth in dragons with temperature induced sex-reversal (Pogona vitticeps, Lacertilia: Agamidae)” by Daniel Hoops, Annalise D. Ceddia, Richard Peters, Sarah Whiteley, Shaun P. Collin, and Jenna M. Crowe-Riddell. Each figure and table included below is cited in the main manuscript.

### Supplementary Figure 1

These figures illustrate the linear measurements that were included in our shape analyses.

### Supplementary Figure 2

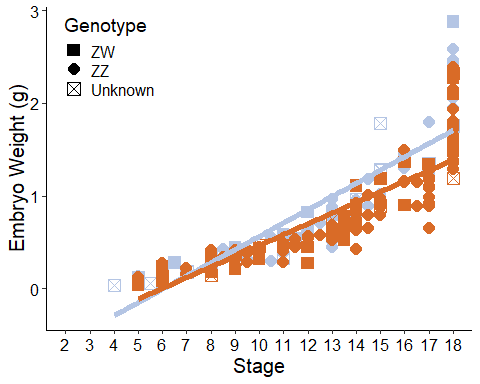

This graph illustrates that embryos incubated at a relatively warm temperature weigh less than embryos incubated at a normal (relatively cool) temperature. This is a replication of the analysis presented in Figure 1c in the main manuscript. This replication uses the complete dataset of embryo weights from Whiteley et al. (2017).

Whiteley SL, Holleley CE, Ruscoe WA, Castelli M, Whitehead DL, Lei J, et al. (2017): Sex determination mode does not affect body or genital development of the central bearded dragon (Pogona vitticeps). EvoDevo 8:25.

Summary Statistics:

| Temp.char | Genotype | N | WTL | sd | se | ci |
| --- | --- | --- | --- | --- | --- | --- |
| 28 °C | ZW | 19 | 0.80 | 0.73 | 0.17 | 0.35 |
| 28 °C | ZZ | 46 | 0.81 | 0.66 | 0.10 | 0.19 |
| 28 °C | Unknown | 17 | 0.60 | 0.64 | 0.16 | 0.33 |
| 36 °C | ZW | 32 | 0.77 | 0.66 | 0.12 | 0.24 |
| 36 °C | ZZ | 130 | 0.66 | 0.50 | 0.04 | 0.09 |
| 36 °C | Unknown | 10 | 0.77 | 0.67 | 0.21 | 0.48 |

A linear model revealed Significant differences in the weight of the embryos between temperature treatments and stages.

```
##
## Call:
## lm(formula = Emb.w.lncs ~ Stage.cs + Genotype + Temp, data = data2017[data2017$Temp.char !=
##     "NA °C" & data2017$Genotype != "Unknown", ])
##
## Residuals:
##      Min       1Q   Median       3Q      Max
## -1.60304 -0.14649  0.02603  0.17587  0.61774
##
## Coefficients:
##              Estimate Std. Error t value Pr(>|t|)
## (Intercept)  0.785184   0.172215   4.559 8.46e-06 ***
## Stage.cs     0.994609   0.020472  48.585  < 2e-16 ***
## GenotypeZZ   0.035927   0.044047   0.816    0.416
## Temp        -0.026396   0.005084  -5.192 4.67e-07 ***
## ---
## Signif. codes:  0 '***' 0.001 '**' 0.01 '*' 0.05 '.' 0.1 ' ' 1
##
## Residual standard error: 0.2755 on 223 degrees of freedom
##   (3 observations deleted due to missingness)
## Multiple R-squared:  0.9143, Adjusted R-squared:  0.9132
## F-statistic: 793.5 on 3 and 223 DF,  p-value: < 2.2e-16
```

### Supplementary Figure 3

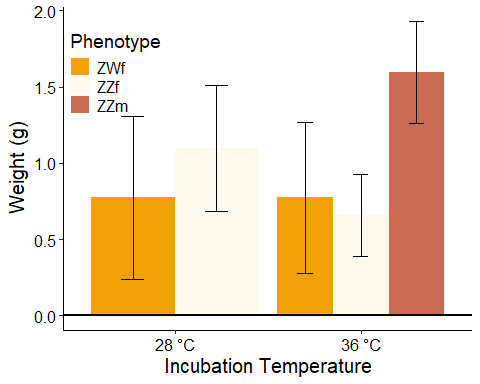

This graph illustrates the weights of embryos with respect to incubation temperature and phenotype. Embryos of unknown phenotype are excluded from the analysis.

Summary Statistics:

| Phenotype | Temp.char | N | Embryo.weight | sd | se | ci | upper.CL | lower.CL |
| --- | --- | --- | --- | --- | --- | --- | --- | --- |
| ZWf | 28 °C | 8 | 0.77 | 0.77 | 0.27 | 0.64 | 1.31 | 0.24 |
| ZWf | 36 °C | 9 | 0.78 | 0.76 | 0.25 | 0.58 | 1.27 | 0.28 |
| ZZf | 28 °C | 12 | 1.10 | 0.73 | 0.21 | 0.47 | 1.51 | 0.68 |
| ZZf | 36 °C | 2 | 0.66 | 0.20 | 0.14 | 1.75 | 0.93 | 0.39 |
| ZZm | 36 °C | 9 | 1.59 | 0.51 | 0.17 | 0.39 | 1.93 | 1.26 |
| Unknown | 28 °C | 6 | 0.23 | 0.19 | 0.08 | 0.20 | 0.38 | 0.07 |
| Unknown | 36 °C | 11 | 0.32 | 0.11 | 0.03 | 0.07 | 0.38 | 0.25 |

A linear model reveals no significant differences between phenotypes in embryo weight, however there are significant effects of stage and incubation temperature.

```
##
## Call:
## lm(formula = Emb.w.lncs ~ Stage.cs + Phenotype + Incubation.temperature,
##     data = data[data$Phenotype != "Unknown", ])
##
## Residuals:
##      Min       1Q   Median       3Q      Max
## -0.95354 -0.08716  0.01214  0.11974  0.39812
##
## Coefficients:
##                        Estimate Std. Error t value Pr(>|t|)
## (Intercept)             1.29114    0.41785   3.090  0.00391 **
## Stage.cs                0.93784    0.04860  19.297  < 2e-16 ***
## PhenotypeZZf           -0.02961    0.10142  -0.292  0.77201
## PhenotypeZZm            0.08433    0.12667   0.666  0.50992
## Incubation.temperature -0.04083    0.01281  -3.186  0.00303 **
## ---
## Signif. codes:  0 '***' 0.001 '**' 0.01 '*' 0.05 '.' 0.1 ' ' 1
##
## Residual standard error: 0.2492 on 35 degrees of freedom
## Multiple R-squared:  0.9335, Adjusted R-squared:  0.9259
## F-statistic: 122.9 on 4 and 35 DF,  p-value: < 2.2e-16
```

Although the graph suggests that, within the embryos incubated at 36C, the ZZm embryos may be larger than the ZWf and ZZf embryos, a posthoc linear model looking at just the 36C embryos with phenotype and stage as predictors does not bear this out.

```
##
## Call:
## lm(formula = Emb.w.lncs ~ Stage.cs + Phenotype, data = data[data$Phenotype !=
##     "Unknown" & data$Incubation.temperature == 36, ])
##
## Residuals:
##      Min       1Q   Median       3Q      Max
## -0.61039 -0.10312  0.07249  0.12912  0.46366
##
## Coefficients:
##              Estimate Std. Error t value Pr(>|t|)
## (Intercept)  -0.21027    0.08617  -2.440   0.0267 *
## Stage.cs      1.14057    0.08668  13.159 5.35e-10 ***
## PhenotypeZZf  0.04189    0.20335   0.206   0.8394
## PhenotypeZZm -0.11874    0.15553  -0.763   0.4563
## ---
## Signif. codes:  0 '***' 0.001 '**' 0.01 '*' 0.05 '.' 0.1 ' ' 1
##
## Residual standard error: 0.2583 on 16 degrees of freedom
## Multiple R-squared:  0.9429, Adjusted R-squared:  0.9322
## F-statistic: 88.13 on 3 and 16 DF,  p-value: 3.658e-10
```

### Supplementary Figure 4

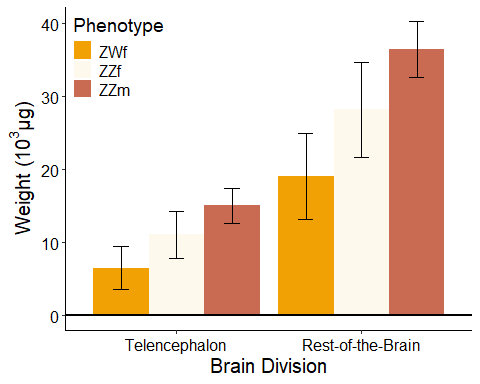

This graph illustrates the size of the telencephalon and the rest-of-the-brain with respect to embryo phenotype.

Summary Statistics:

| Phenotype | N | Weight | sd | se | ci | Division | upper.CL | lower.CL |
| --- | --- | --- | --- | --- | --- | --- | --- | --- |
| ZWf | 17 | 6.55 | 6.22 | 1.51 | 3.20 | Telencephalon | 9.51 | 3.59 |
| ZZf | 14 | 11.08 | 6.21 | 1.66 | 3.59 | Telencephalon | 14.33 | 7.83 |
| ZZm | 9 | 15.04 | 3.62 | 1.21 | 2.79 | Telencephalon | 17.41 | 12.68 |
| Unknown | 17 | 3.45 | 2.39 | 0.58 | 1.23 | Telencephalon | 4.59 | 2.32 |
| ZWf | 17 | 19.14 | 12.37 | 3.00 | 6.36 | Rest-of-the-Brain | 25.02 | 13.26 |
| ZZf | 14 | 28.24 | 12.39 | 3.31 | 7.15 | Rest-of-the-Brain | 34.73 | 21.75 |
| ZZm | 9 | 36.47 | 5.80 | 1.93 | 4.46 | Rest-of-the-Brain | 40.26 | 32.67 |
| Unknown | 17 | 13.71 | 8.08 | 1.96 | 4.15 | Rest-of-the-Brain | 17.55 | 9.87 |

Two ANOVAs, one with telencephalon weight as the response variable and one as the rest of the brain weight, reveal that, for both divisions, there are significant effects with respect to stage but not phenotype. Incubation temperature is significant with respect to the telencephalon but not the rest of the brain.

```
## [1] "Telencephalon:"
## Anova Table (Type II tests)
##
## Response: Emb.w.lncs
##                         Sum Sq Df  F value  Pr(>F)
## Stage.cs               23.1259  1 372.3727 < 2e-16 ***
## Phenotype               0.0413  2   0.3327 0.71924
## Incubation.temperature  0.6304  1  10.1508 0.00303 **
## Residuals               2.1736 35
## ---
## Signif. codes:  0 '***' 0.001 '**' 0.01 '*' 0.05 '.' 0.1 ' ' 1
## [1] "Rest-of-the-Brain:"
## Anova Table (Type II tests)
##
## Response: Rest.w.lncs
##                         Sum Sq Df F value    Pr(>F)
## Stage.cs               17.0024  1 81.9149 1.078e-10 ***
## Phenotype               0.1063  2  0.2561    0.7755
## Incubation.temperature  0.0875  1  0.4217    0.5204
## Residuals               7.2647 35
## ---
## Signif. codes:  0 '***' 0.001 '**' 0.01 '*' 0.05 '.' 0.1 ' ' 1
```

### Supplementary Figure 5

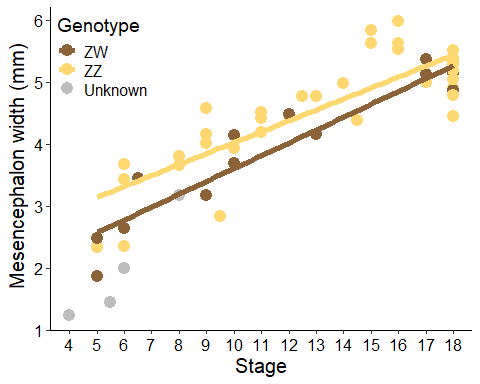

This figure illustrates the width of the telencephalon with respect to genotype and incubation temperature. Because the widest part of the mesencephalon at all developmental stages examined is the optic tectum, this is effectively an analysis of the width of the optic tectum.

Summary Statistics:

| Temp.char | Genotype | N | MWM | sd | se | ci | upper.CL | lower.CL |
| --- | --- | --- | --- | --- | --- | --- | --- | --- |
| 28 °C | ZW | 7 | 3.91 | 1.13 | 0.43 | 1.04 | 4.75 | 3.08 |
| 28 °C | ZZ | 12 | 4.69 | 0.97 | 0.28 | 0.62 | 5.24 | 4.14 |
| 36 °C | ZW | 8 | 4.05 | 1.12 | 0.39 | 0.93 | 4.82 | 3.28 |
| 36 °C | ZZ | 22 | 4.48 | 0.91 | 0.19 | 0.41 | 4.86 | 4.10 |

MWM = Mesencephalon width

A linear model reveals a significant effect of stage and genotype, but not incubation temperature, on mesencephalon width (effectively optic tectum width because the optic tectum is the largest part of the mesencephalon).

```
##
## Call:
## lm(formula = MWM.lncs ~ Stage + Genotype + Temp.char, data = data[data$Genotype !=
##     "Unknown", ])
##
## Residuals:
##     Min      1Q  Median      3Q     Max
## -1.0655 -0.1967  0.1163  0.2660  0.5780
##
## Coefficients:
##                Estimate Std. Error t value Pr(>|t|)
## (Intercept)    -1.68926    0.19164  -8.815 2.32e-11 ***
## Stage           0.13679    0.01276  10.724 5.57e-14 ***
## GenotypeZZ      0.27563    0.12427   2.218   0.0316 *
## Temp.char36 °C -0.08328    0.11677  -0.713   0.4794
## ---
## Signif. codes:  0 '***' 0.001 '**' 0.01 '*' 0.05 '.' 0.1 ' ' 1
##
## Residual standard error: 0.3959 on 45 degrees of freedom
##   (4 observations deleted due to missingness)
## Multiple R-squared:  0.7386, Adjusted R-squared:  0.7212
## F-statistic: 42.39 on 3 and 45 DF,  p-value: 3.644e-13
```

### Supplementary Figure 6

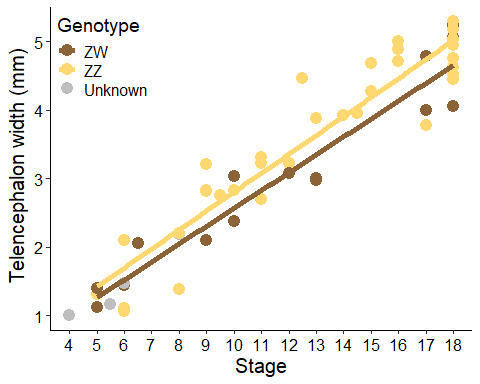

This figure illustrates the width of the telencephalon with respect to genotype and stage.

Summary Statistics:

| Temp.char | Genotype | N | MWT | sd | se | ci | upper.CL | lower.CL |
| --- | --- | --- | --- | --- | --- | --- | --- | --- |
| 28 °C | ZW | 7 | 2.99 | 1.53 | 0.58 | 1.42 | 4.12 | 1.85 |
| 28 °C | ZZ | 12 | 3.96 | 1.32 | 0.38 | 0.84 | 4.71 | 3.22 |
| 36 °C | ZW | 8 | 3.09 | 1.25 | 0.44 | 1.05 | 3.96 | 2.23 |
| 36 °C | ZZ | 22 | 3.44 | 1.25 | 0.27 | 0.55 | 3.97 | 2.92 |

MTW = Telencephalon width

A linear model reveals a significant effect of stage, but not genotype or incubation temperature, on telencephalon width.

```
##
## Call:
## lm(formula = MWT.lncs ~ Stage + Genotype + Temp.char, data = data[data$Genotype !=
##     "Unknown", ])
##
## Residuals:
##      Min       1Q   Median       3Q      Max
## -1.09051 -0.22636  0.03555  0.26550  0.54753
##
## Coefficients:
##                Estimate Std. Error t value Pr(>|t|)
## (Intercept)    -2.24782    0.17527 -12.825   <2e-16 ***
## Stage           0.18604    0.01167  15.947   <2e-16 ***
## GenotypeZZ      0.18787    0.11365   1.653   0.1053
## Temp.char36 °C -0.18121    0.10680  -1.697   0.0967 .
## ---
## Signif. codes:  0 '***' 0.001 '**' 0.01 '*' 0.05 '.' 0.1 ' ' 1
##
## Residual standard error: 0.3621 on 45 degrees of freedom
##   (4 observations deleted due to missingness)
## Multiple R-squared:  0.8561, Adjusted R-squared:  0.8465
## F-statistic: 89.25 on 3 and 45 DF,  p-value: < 2.2e-16
```

### Supplementary Figure 7

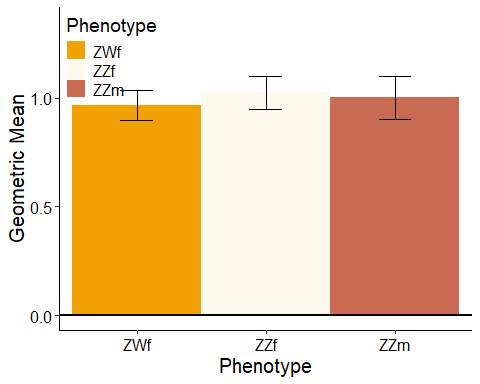

This figure illustrates the brain geometric mean (a measure of brain size) with respect to phenotype.

Summary Statistics:

| Phenotype | Temp.char | N | geomean | sd | se | ci |
| --- | --- | --- | --- | --- | --- | --- |
| ZWf | 28 °C | 6 | 0.89 | 0.24 | 0.10 | 0.26 |
| ZWf | 36 °C | 8 | 0.87 | 0.31 | 0.11 | 0.26 |
| ZZf | 28 °C | 11 | 1.04 | 0.18 | 0.06 | 0.12 |
| ZZf | 36 °C | 2 | 1.04 | 0.00 | 0.00 | 0.01 |
| ZZm | 36 °C | 9 | 1.11 | 0.05 | 0.02 | 0.04 |
| Unknown | 28 °C | 6 | 0.52 | 0.38 | 0.16 | 0.40 |
| Unknown | 36 °C | 11 | 0.90 | 0.15 | 0.05 | 0.10 |

An ANOVA reveals a significant effect of stage, but not phenotype incubation temperature, on brain geometric mean.

```
## Anova Table (Type II tests)
##
## Response: geomean
##            Sum Sq Df F value    Pr(>F)
## Stage     0.90165  1 66.2919 3.396e-09 ***
## Phenotype 0.01864  2  0.6851    0.5115
## Temp.char 0.01845  1  1.3562    0.2531
## Residuals 0.42164 31
## ---
## Signif. codes:  0 '***' 0.001 '**' 0.01 '*' 0.05 '.' 0.1 ' ' 1
```

### Supplementary Table 1

```
##
## Analysis of Variance, using Residual Randomization
## Permutation procedure: Randomization of null model residuals
## Number of permutations: 1000
## Estimation method: Ordinary Least Squares
## Sums of Squares and Cross-products: Type I
## Effect sizes (Z) based on F distributions
##
##           Df      SS      MS     Rsq       F      Z Pr(>F)
## Stage      1 10.3912 10.3912 0.72566 94.8442 5.5119  0.001 ***
## Phenotype  2  0.3315  0.1658 0.02315  1.5129 0.9608  0.185
## Temp       1  0.2005  0.2005 0.01400  1.8302 1.0972  0.136
## Residuals 31  3.3964  0.1096 0.23718
## Total     35 14.3196
## ---
## Signif. codes:  0 '***' 0.001 '**' 0.01 '*' 0.05 '.' 0.1 ' ' 1
##
## Call: procD.lm(f1 = lsr ~ Stage + Phenotype +
## Temp, data = analyze_lsr_phen)
```

This permutational ANOVA demonstrates that there is an effect of stage, but not incubation temperature or phenotype, on brain shape.

### Supplementary Table 2

```
##
## Analysis of Variance, using Residual Randomization
## Permutation procedure: Randomization of null model residuals
## Number of permutations: 1000
## Estimation method: Ordinary Least Squares
## Sums of Squares and Cross-products: Type I
## Effect sizes (Z) based on F distributions
##
##           Df     SS      MS     Rsq      F       Z Pr(>F)
## Stage      1 1.8299 1.82987 0.34152 8.0786  3.5192  0.001 ***
## Genotype   1 0.1268 0.12680 0.02367 0.5598 -0.4511  0.671
## Temp       1 0.2303 0.23029 0.04298 1.0167  0.3023  0.353
## Residuals 14 3.1711 0.22651 0.59184
## Total     17 5.3581
## ---
## Signif. codes:  0 '***' 0.001 '**' 0.01 '*' 0.05 '.' 0.1 ' ' 1
##
## Call: procD.lm(f1 = lsr ~ Stage + Genotype +
## Temp, data = analyze_lsr_early)
```

This permutational ANOVA demonstrates that there is a significant effect of stage, but not genotype or incubation temperature, on the shapes of embryo brains between stages 4 and 10 (i.e. “early” developmental stages). ## Supplementary Table 3

```
##
## Analysis of Variance, using Residual Randomization
## Permutation procedure: Randomization of null model residuals
## Number of permutations: 1000
## Estimation method: Ordinary Least Squares
## Sums of Squares and Cross-products: Type I
## Effect sizes (Z) based on F distributions
##
##           Df     SS      MS     Rsq       F       Z Pr(>F)
## Stage      1 2.4851 2.48506 0.52857 36.8043  3.7313  0.001 ***
## Genotype   1 0.0444 0.04437 0.00944  0.6572 -0.0462  0.519
## Temp       1 0.3490 0.34899 0.07423  5.1686  2.2465  0.009 **
## Residuals 27 1.8231 0.06752 0.38776
## Total     30 4.7015
## ---
## Signif. codes:  0 '***' 0.001 '**' 0.01 '*' 0.05 '.' 0.1 ' ' 1
##
## Call: procD.lm(f1 = lsr ~ Stage + Genotype +
## Temp, data = analyze_lsr_late)
```

This permutational ANOVA demonstrates that there is are significant effects of stage and incubation temperature, but not genotype, on the shapes of embryo brains between stages 11 and 18 (i.e. “late” developmental stages).
